# A minor impact of VGLUT1 expression level on quantal size revealed through the characterization of VGLUT1^mEos2^ knock-down new mouse model

**DOI:** 10.1101/2024.09.23.614439

**Authors:** Xiao Min Zhang, Urielle François, Maria Florencia Angelo, Stéphane Claverol, Magali Mondin, Christelle Martin, Melissa Deshors, Yann Humeau, Noa Lipstein, Etienne Herzog

## Abstract

Synaptic vesicles (SVs) are small organelles secreting neurotransmitters at synapses. By fusing a photoactivated fluorescent protein to VGLUT1, we generated a VGLUT1^mEos2^ knock-in mouse. VGLUT1^mEos2^ knock-in mice are viable and healthy, but exhibit a severe reduction in VGLUT1 expression levels. Using VGLUT1^mEos2^ expressing neurons, we established paradigms to trace individual SV mobility at the single-molecule level or via massive photoconversion. Hippocampal neurons with significantly diminished VGLUT1 expression maintain unaltered miniature glutamate release characteristics in terms of quantal size and frequency. We demonstrate that VGLUT1 expression level are not correlated in a linear fashion with the vesicular glutamate content. In conclusion, the VGLUT1^mEos2^ mouse line serves as a powerful tool for exploring SV mobility properties and elucidating the contributions of VGLUT1 to excitatory neurotransmission and cognitive processes.

## Introduction

Synaptic vesicles (SVs) are strategically clustered at the pre-synaptic terminals of neurons. These vesicles play a pivotal role in neurotransmission by releasing their neurotransmitter content to initiate the transfer of information from one neuron to another. Despite the extensive research on the SV cycle, release probability, and the mechanisms of neurotransmitter loading into the SV, comprehensive insights into SV mobility and quantal heterogeneity remain elusive.

Synapses have long been thought to represent autonomous units relying on local recycling and delivery of de novo synthesized components delivered from the cell soma through axonal transport (Bonanomi et al., 2006). More and more data suggest that neurite-wide exchange of material is occurring and is necessary for proper network function (Darcy et al., 2006; Kalla et al., 2006; Tsuriel et al., 2006; Westphal et al., 2008; Minerbi et al., 2009; Herzog et al., 2011). The analysis of SV exchange between *en passant* boutons has been most extensively studied, leading to the notion of an SV super-pool that spans the axon or a segment of it. This SV super-pool was defined through morphological observations, contrasting with the classical SV pools, that were identified based on functional properties in electrophysiological paradigms (Denker and Rizzoli, 2010).

Glutamate is the most abundant excitatory neurotransmitter. Vesicular GLUtamate Transporter 1 (VGLUT1) is the main protein responsible for loading glutamate into SVs in cortical and hippocampal neurons. Fluorescence labeling of VGLUT1 enables tracking of glutamatergic SVs’ in axons. In our previous study, we successfully observed SV trafficking at cortical synapses in living mice using a VGLUT1^venus^ knock-in (KI) mouse line, that tags the endogenous VGLUT1 with a fluorescent reporter protein VENUS (Herzog et al., 2011). These mice were further used to establish the physical determinants of SV mobility at cerebellar mossy fiber terminals under different activity regimes (Rothman et al., 2016). In addition, our prior research has demonstrated that the structure of VGLUT1 affects SVs mobility and neurotransmission through its interaction with endophilin A1 (Siksou et al., 2013; Zhang et al., 2019). Thus, using bulk imaging approaches such as FRAP and time-lapse imaging, we have been able to track SV mobility either *in vitro* or *in vivo*. However, these measurements lacked the capability to resolve the motion of individual synaptic vesicles in living neurons. Resolving this would provide deeper insights into how SV mobility, turnover at synapses and exchange with the super-pool contribute to neurotransmission and synaptic plasticity.

The quantal postsynaptic currents result from the release of neurotransmitter by a single vesicle activating the pool of ionotropic receptors positioned in the synapse. The size of this quantal response is dictated by several parameters including the quantal content - the amount of neurotransmitter within the vesicle. Adjusting this quantal size is critical for diverse forms of synaptic plasticity (Edwards, 2007; Sakaba et al., 2002; Turrigiano, 2012) and SV release probability (Walmsley et al., 1988). Experimental evidence has shown that altering vesicular transporter levels can significantly affect quantal size (Takamori, 2016). Overexpression of transporters for acetylcholine and dopamine leads to increased quantal sizes (Song et al., 1997; Lohr et al., 2014). Conversely, reduced expression of vesicular monoamine transporter 2 (VMAT2) diminishes the storage capacity for dopamine and catecholamines (Caudle et al., 2007; Taylor et al., 2014). Similarly, knockdown of VGLUT2 reduces miniature excitatory postsynaptic current (mEPSC) amplitudes in thalamic neurons (Moechars et al., 2006).

Research has indicated that VGLUT1 expression directly governs the glutamate content of SVs, with its expression level directly correlating to the vesicular glutamate content: absence of VGLUT1 results in decreased content, while overexpression increases the SV’s quantal size (Fremeau et al., 2004; Wojcik et al., 2004). However, research conducted in Drosophila has challenged this linear relationship, revealing that even one copy of dVGLUT was sufficient to load wild-type (WT) levels of glutamate (Daniels et al., 2006). Therefore, developing a more precise method to control VGLUT1 expression levels would greatly facilitate understanding the specific impact of VGLUT1 on SV glutamate loading and, by extension, synaptic transmission strength.

Here, we report a new VGLUT1^mEos2^ KI line. VGLUT1 is fused to a photo-switchable fluorescent protein mEos2, which is widely used for pointillist super-resolution techniques (McKinney et al., 2009; Nienhaus et al., 2005). VGLUT1^mEos2^ exhibited the anticipated normal distribution and photoactivatable properties, but also a significant reduction in VGLUT1 expression level. Yet, our findings indicate that VGLUT1^mEos2^ heterozygote mice provide a good model for the investigations on SV mobility. Moreover, using both VGLUT1^mEos2^ mice and VGLUT1 knockout (KO) mice, we were able to establish that the expression level of VGLUT1 does not correlate with a linear change in quantal size.

## Results

### Tracing SVs mobility with VGLUT1^mEos2^

Photoactivatable fluorescent proteins, such as mEos2, have been extensively employed in super-resolution imaging to pinpoint and track single molecules (Deschout et al., 2014; Shroff et al., 2008; Subach et al., 2009). This green-to-red photoconvertible protein allows for the real-time monitoring of targeted protein localization within live cells (McKinney et al., 2009). Here, we engineered a construct by fusing mEos2 to the C-terminus of VGLUT1, as previously done with VENUS (Herzog et al., 2011). VGLUT1^mEos2^ should allow to investigate the mobility of single SV.

As a proof-of-concept experiment, we over-expressed VGLUT1^mEos2^ in dissociated primary hippocampal neuron cultures using lentivirus transduction. Under 488 nm laser illumination, we observed presynaptic boutons filled with VGLUT1^mEos2^ emitting green fluorescence as expected (Fig. 1A) (Herzog et al., 2011). To achieve single-molecule imaging with a resolution of 20-40 nm, we applied a moderate 405 nm laser excitation (3 W/cm²) to photoconvert the fluorescence. The red VGLUT1^mEos2^ signals were then captured following excitation at 561 nm.

**Figure 1.**
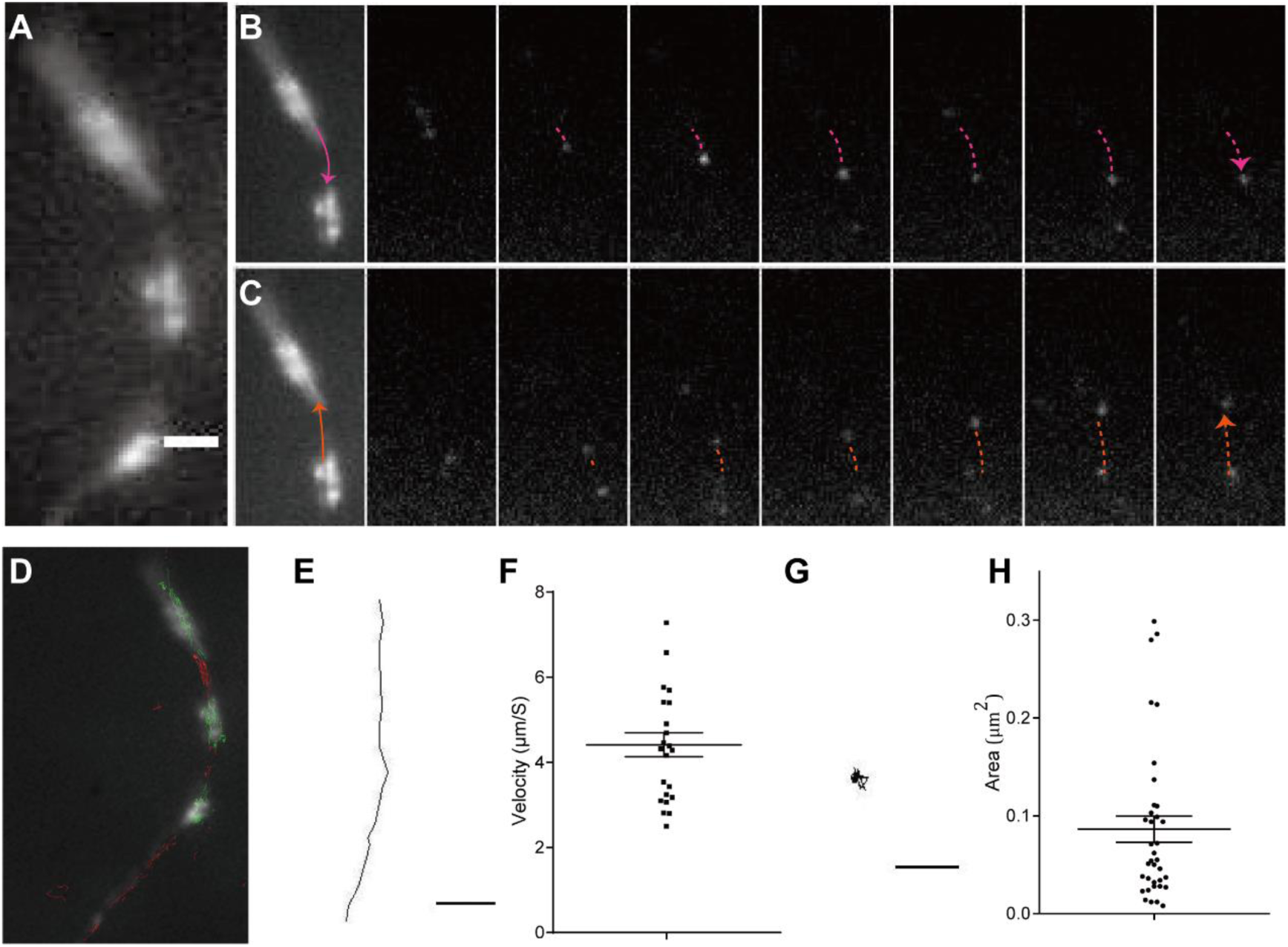
PALM imaging with VGLUT1^mEos2^ to investigate synaptic vesicle mobility. (A) Expression of VGLUT1 fused to mEos2 (VGLUT1^mEos2^) in hippocampal neurons at 18 days in culture. Scale bar 1 *µm*. (B & C) Examples of single synaptic vesicles moving in both directions between synaptic boutons of the same axon. (D) Traces recorded outside of the bouton (in red) and inside the bouton (in green) were overlayed with the axon. (E) A single trace of SV moving outside of boutons. (F) The speed of SV direct motion. (G) A single trace of SV confined inside the bouton. (H) The area of confined traces within boutons. For (F), (H), data are represented as mean ± SEM

As VGLUT1 is highly selectively and efficiently addressed to SVs with a minimal pool of the protein on other subcellular compartments (Foss et al., 2013; Pan et al., 2015), one can assume that tracking individual VGLUT1^mEos2^ proteins matches tracking the mobility of single SVs. Our imaging revealed that individual SVs could be traced moving along axons both anterogradely and retrogradely (Fig. 1B & C). Upon overlaying these movement traces onto the initial 488 nm images (Fig. 1A), we categorized them into two distinct groups: those confined within synaptic boutons and those exhibiting linear trajectories outside the boutons (Fig. 1D). We focused on the representative straight-line traces, indicative of uninterrupted and unidirectional SV movement along the axon (Fig. 1E). Analyzing these traces allowed us to estimate the instantaneous speed of SVs in the axons, which varied between 2.50 and 7.28 µm/s, with an average speed of 4.32 ± 1.30 µm/s (Fig. 1F). Subsequently, we delved into the diffusion coefficient (*D*) characteristics of SV traces within synaptic boutons (Fig. 1G) through a classical mean square displacement (MSD) analysis (Penn et al., 2017). At this stage, our data reveal that the majority of SVs displayed confined movement with an area of diffusion less than 0.1 µm² (Fig. 1H). Establishing a direct correlation between each observed diffusion pattern and a specific stage in the SV life cycle will require further investigations. Altogether, these experiments validate the expression and trafficking of VGLUT1^mEos2^ fusion proteins in view of the generation of a knock-in model.

### Generation of VGLUT1^mEos2^ knock-in mouse

Given the successful demonstration of single SV tracing using VGLUT1^mEos2^ cDNA constructs, we proceeded to create a VGLUT1^mEos2^ KI mouse line, which would enable the monitoring of endogenous, rather than overexpressed VGLUT1. We adopted the same strategy as previously employed in our laboratory for generating the VGLUT1^Venus^ KI mouse (Herzog et al., 2011). This involved fusing mEos2 cDNA to exon 12 of the Slc17a7 gene (encoding the VGLUT1 protein) before the endogenous stop codon and incorporating a neomycin resistance cassette into the 3’ untranslated region (Fig. 2A). Homologous recombination in embryonic stem cells was carried out followed by injection into blastocytes. With this we could generate offspring harboring the VGLUT1^mEos2Neo^ allele (Fig. 2A). To mitigate any potential detrimental effects on VGLUT1 expression caused by the neomycin resistance cassette, VGLUT1^mEos2Neo/+^ mice were crossed with EIIa-cre mice, which carry the cre transgene under the adenovirus EIIa promoter, ensuring Cre recombinase expression during early embryonic development (Lakso et al., 1996). The offspring from this cross ultimately led to the generation of homozygous mice without the neomycin resistance cassette (VGLUT1^Eo/Eo^; Fig. 2A).

**Figure 2.**
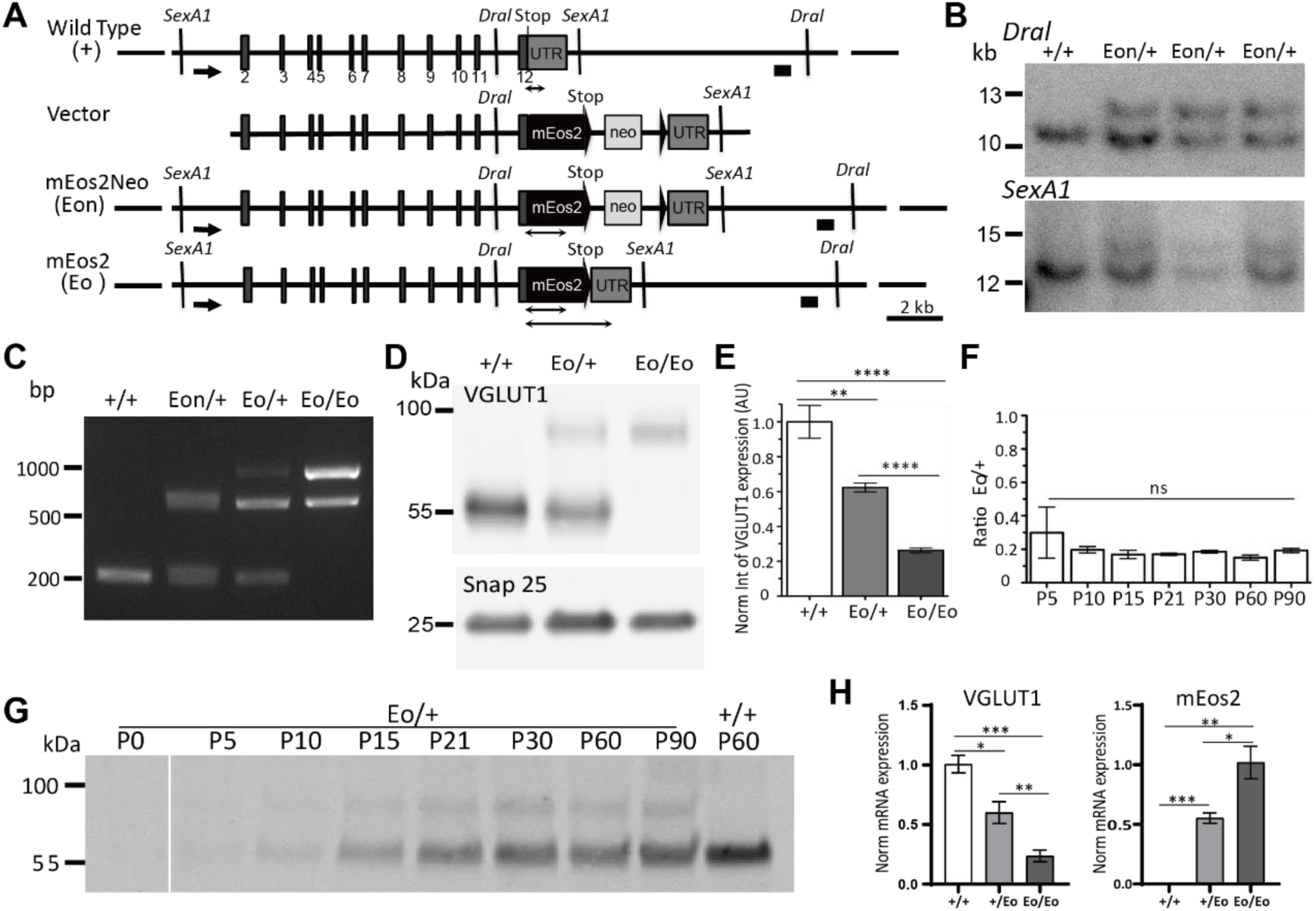
Generation of VGLUT1^mEos2^ KI mice. (A) Strategy for the generation of the VGLUT1^mEos2^ KI mutation in mouse embryonic stem cells. The WT VGLUT1 gene, targeting vector, mutated gene after homologous recombination (mEos2Neo, Eon), and mutated gene after Cre recombination (mEos2, Eo) are shown. Exons are indicated by gray boxes. The black triangles indicate loxP sites. Two groups of black horizontal bars (solid line for DraI restriction, dash line with arrow for SexA1 restriction) indicate the 5’ and 3’ probes used for Southern analysis. Double arrows represent the amplicons of PCR genotyping in the various genotypes. NEO, Neomycin resistance gene; mEos2, the open reading frame of mEos2 fluorescent protein; UTR, 3’ untranslated region. (B) Southern blot analysis of ES cell DNA after DraI and SexA1 restriction using the probe indicated in A. Eon, mutated allele with neomycin resistance cassette (12.7 kb for DraI, 13.9 kb for SexA1); +, WT allele (10.8 kb for DraI,12.1 kb for SexA1). (C) PCR genotyping of WT, mEos2Neo, and mEos2 mouse tail DNA. (D) Comparison of VGLUT1 and VGLUT1^mEos2^ expression in VGLUT1^mEos2^ mouse line. mEos2 tagged VGLUT1 is ∼25 kDa larger than the WT transporter. (E) Semiquantitative western blot analysis of brain homogenates from +/+ (n = 3), Eo/+ (n = 6), and Eo/Eo (n = 5). VGLUT1 signals were normalized to SNAP-25 signals. (F) Semiquantitative western blot analyses (near-infrared fluorescent detection) of brain homogenates from Eo/+ at different developmental stages. The ratio between the levels of native and tagged transporters was constant throughout the developmental stages. P, postnatal day. (P5, 0.2995 ± 0.15 AU, n = 2; P10, 0.20 ± 0.02 AU, n = 2; P15, 0.17 ± 0.02 AU, n = 3; P21, 0.17 ± 0.01 AU, n = 3; P30, 0.19 ± 0.01 AU, n = 3; P60, 0.15 ± 0.02 AU, n = 3; P90, 0.19 ± 0.01 AU, n = 3). (G) A representative array of blots was reconstructed from lanes of the same membrane. H. Normalized VGLUT1 and mEos2 mRNA expression levels in +/+, Eo/+, and Eo/Eo animals n = 3 for all genotype animals. For (E), (F), (H), data are represented as mean ± SEM, * for *p* < 0.05, ** for *p* < 0.01, *** for *p* < 0.001, **** for *p* < 0.0001, ns for *p* > 0.05.

For initial screening, we assessed 96 ES cell clones, approximately 70% of which showed the desired homologous recombination as confirmed through Southern blot analysis using both 3’ and 5’ external probes (Fig. 2B). For routine breeding purposes, PCR genotyping was performed (Fig. 2C). VGLUT1^Eo/Eo^ mice were born at the anticipated Mendelian frequency and exhibited no distinguishable differences from their WT and VGLUT1^Eo/+^ littermates. Neither VGLUT1^Eo/+^ nor VGLUT1^Eo/Eo^ mice showed any obvious behavioral or morphological deficits at a cage environment. These animals had a normal life expectancy, were fertile, and did not display any apparent deficits. Although only limited breeding trials were conducted with homozygous VGLUT1^Eo/Eo^ animals, they appeared to be viable and healthy.

To evaluate the expression levels of VGLUT1^mEos2^ in these genetically modified mice, we performed a systematic quantification of VGLUT1 expression in brain homogenates from heterozygous and homozygous animals using western blotting with infrared fluorescence detection. On SDS-PAGE, the mEos2-tagged VGLUT1 protein migrated with the anticipated 25-30 kDa increase compared to its WT counterpart, displaying a marginally more compact migration pattern (Fig. 2D). For accurate comparison, we quantified the normalized integrated intensities of VGLUT1 and VGLUT1^mEos2^ across different genotypes. To ensure consistency in loading and normalization, the SNARE protein SNAP-25 was used as an internal control (Fig. 2D). Unexpectedly, our results revealed a significant reduction in VGLUT1^mEos2^ expression (Fig. 2E). This knockdown effect was consistent for both VGLUT1^Eo/+^ and VGLUT1^Eo/Eo^ mice, indicating that the presence of the VGLUT1^mEos2^ construct led to a decrease in VGLUT1 expression to approximately one-fourth of the WT level.

We next monitored the developmental pattern of WT and tagged VGLUT1 expression in VGLUT1^Eo/+^ mice. We conducted a series of western blot analyses on brain homogenates from mice ranging in age from postnatal day P0 to P90 (Fig. 2F). The ratio between the WT and fusion protein remained constant throughout postnatal development, with VGLUT1^mEos2^ expression consistently representing about one-fourth of the WT VGLUT1 (Fig. 2F and 2G). To discern the mechanism of mEos2 fusion knock-down we proceeded with quantitative RT-PCR (RT-qPCR) assays to evaluate VGLUT1 and VGLUT1^mEos2^ mRNA expressions across different genotypes. VGLUT1^mEos2^ mRNA expression was consistently reduced compared to the WT VGLUT1 expression with a ratio equivalent to the one observed for protein levels (Fig. 2H). We conclude that the inclusion of the mEos2 cDNA in the Slc17a7 gene sequence results in lower mRNA levels, consequently leading to lower VGLUT1^mEos2^ protein levels.

### Localization of mEos2 fluorescence in VGLUT1^mEos2^ mouse brain

As anticipated, upon fixation with paraformaldehyde (PFA), VGLUT1^Eo/Eo^ brain tissue displayed intense green fluorescence when illuminated with blue light (488 nm). In sagittal sections of the brain, significant VGLUT1^mEos2^ fluorescence was observed in regions known for dense VGLUT1 innervation, including the olfactory bulb, cortex, striatum, hippocampus, thalamus, and cerebellar cortex (Fig. 3A). Upon closer examination at higher magnifications, the characteristic VGLUT1 labeling patterns were evident within the neuropil of the hippocampus, cortex, and cerebellum (Fig. 3B-D), while no mEos2 signal was detected in cell body layers, confirming that mEos2 tagging did not disrupt the qualitative distribution of VGLUT1, which is the same as seen with the VGLUT1^Venus^ mouse brains (Herzog et al., 2011).

**Figure 3.**
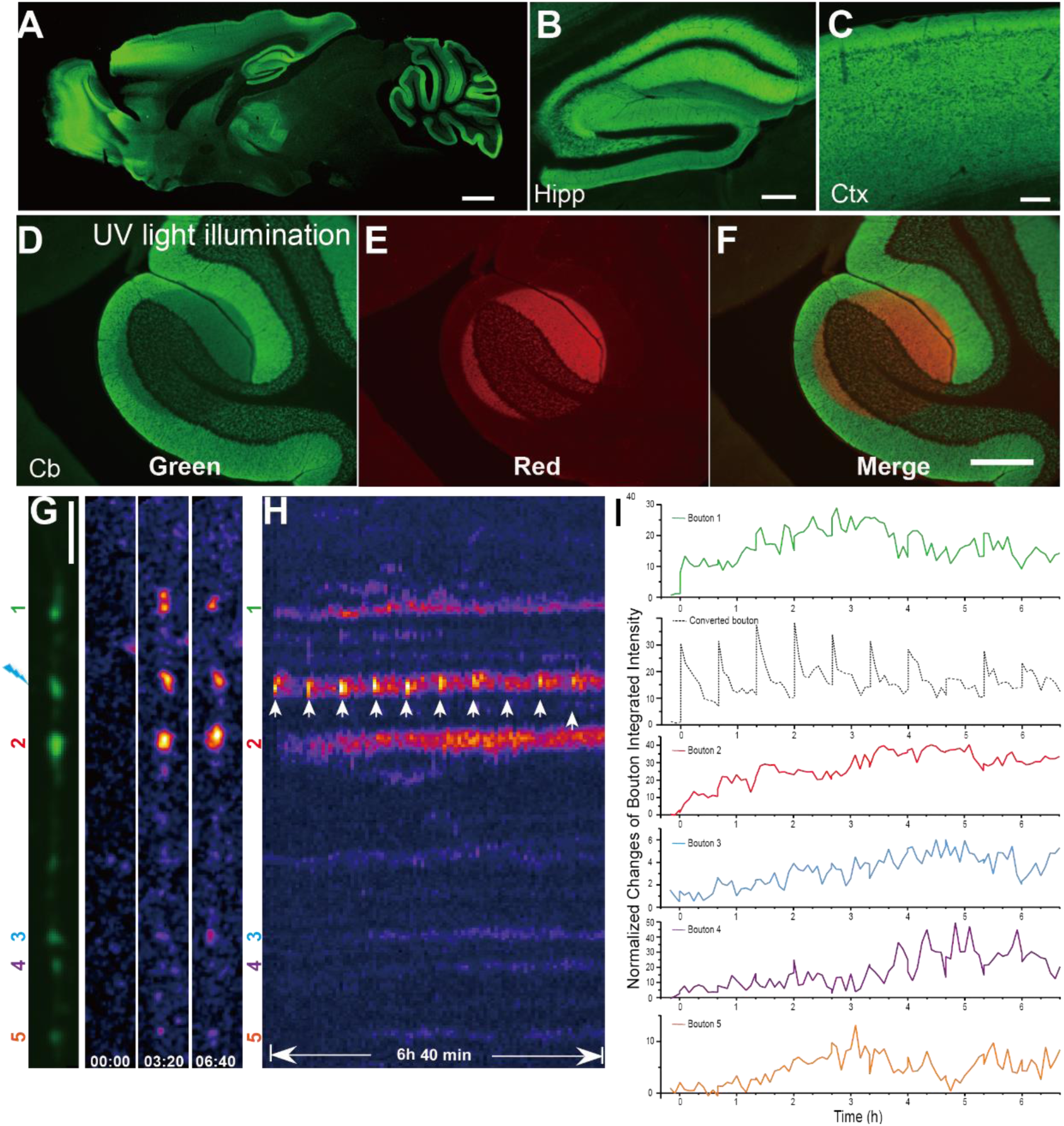
Localization, photoconversion and dynamics of VGLUT1^mEos2^ fluorescence. (A) Overview of direct VGLUT1^mEos2^ fluorescence sagittal section of Eo/Eo brain. Scale bar, 1 mm. (B, C & D) Higher magnification images depicting hippocampus (Hipp), cortex (Cx), and cerebellum (Cb). Fine puncta of fluorescent material are concentrated in neuropil areas of conventional VGLUT1 positive regions. Scale bar in B & C, 200 µm. (D, E & F) Test photoconversion of a big spot in the cerebellum. Scale bar in F, 300 µm. (G & H) Imaging SV super-pool with VGLUT1^Eo/+^ culture. An axonal fiber with 6 *en passant* boutons was selected. One synapse was selected to be converted with a 405 nm laser (blue arrow pointed). G left, neighboring boutons are labeled as 1, 2, 3, 4, and 5 from up to down. Scale bar, 5 µm. G right, the representative image in red channel at time of 0 min, 3 hours 20 min, and 6 hours 40 min. H. Kymographs of the live imaging acquisition in the red channel. The selected bouton was photoconverted with a brief pulse of 405nm excitation and monitored every 5 min for 40 min. The pulses of photoconversion are labeled with white arrows. (I) The intensity plot of each monitored synapse with 561 nm laser excitation.

Upon exposure to a strong ultraviolet light on the brain slice, we observed a clear regional photoconversion, where the green-mEos2 signal converted to red (Fig. 3 D-F). Despite the knock-down of VGLUT1^mEos2^ to ∼25% of WT levels, the mEos2 signal remained strikingly bright, demonstrating its utility for visualizing and tracking VGLUT1 protein even at reduced expression levels.

### Extended SV super-pool monitoring using VGLUT1^mEos2^ photoconversion

We selected VGLUT1^Eo/+^ animals for imaging experiments due to their strong fluorescent signal and moderate reduction in VGLUT1 expression, which provided an ideal compromise. To meticulously examine SV super-pool dynamics, we optimized protocols that involved local mass photoconversion at a chosen synaptic bouton and subsequent tracking of the redistribution of converted red SVs/VGLUT1^mEos2^ molecules.

For this purpose, we established a mixed neuronal culture where VGLUT1^Eo/+^ cells were co-cultured with littermate WT cells at a ratio of 1:10 within the same well. This setup allowed us to easily identify *en passant* boutons aligned along the same axon (Fig. 3G). Despite encountering challenges related to the photophysical properties of mEos2, specifically its tendency to enter a long-lived reversible dark state in the green channel, limiting photoconversion rates and preventing quantitative analysis using the green channel (Berardozzi et al., 2016), we developed an adapted imaging acquisition strategy. Our adapted photoconversion paradigm entailed repeatedly applying photoconversion pulses (405 nm laser, 80 mW) to the same bouton every 40 minutes over ten cycles (Fig. 3H). During each cycle of the imaging process, a snapshot was captured from both the green and red channels at an interval of 5 minutes.

In one illustrative example, we showcased an axon harboring six *en passant* boutons (Fig. 3G). We chose one bouton for recurrent photoconversion to track SV super-pool exchange for several hours along the axon (Fig. 3H, I). The kymograph representation enabled us to trace the temporal changes in red signal distribution and intensity for almost seven hours of continuous imaging.

Upon photoconversion, the targeted bouton exhibited a peak of red fluorescence intensity, followed by gradual dispersion over the next 40 minutes (Fig. 3H). While the photoconverted bouton peaked at each photoconversion cycle, surrounding boutons displayed a slow rise of red signal over time after photoconversion bursts. This behavior is in line with the exchange of photoconverted SVs from the source synapse to the rest of the axon and conversely of green SVs from the axon to the source bouton (Fig. 3H, I) (Herzog et al., 2011; Siksou et al., 2013; Zhang et al., 2019). Notably, immediate neighbors to the source bouton (bouton 1 and 2) didn’t follow the same red fluorescence jumps. Therefore, the photoconversion seems well confined to the target bouton (Fig. 3I). Meanwhile, and in line with previous reports (Bourgeois, 2023), we observed that the 405 nm excitation not only converted the green mEos2 to red mEos2, but also recruited mEos2 from a dark state to the green state (Bourgeois, 2023). Hence, the green intensity cannot report faithfully the quantitative changes induced by SV exchange between boutons (Fig. S1BC). (Darcy et al., 2006; Herzog et al., 2011; Zhang et al., 2019)

Altogether, photoconversion of mEos2 tagged to VGLUT1 allows to highlight the ongoing process of SV exchange along the axon over extended time frames and axonal distance.

### The quantal size of glutamate release is not proportional to VGLUT1 expression level

We could previously show that the tagging of a fluorescent protein on VGLUT1 c-terminus is neutral to the function of VGLUT1 (Herzog et al., 2011). Our collective results in the characterization of the VGLUT1^mEos2^ knock-in (Fig 1-4) strongly advocate that VGLUT1 protein is not disturbed by mEos2 tagging as in the case of tagging with VENUS, rather the induced knock-down is due to a defect at the level of transcription or mRNA stability (Fig 2H). Hence, the severe knock-down of VGLUT1 in the VGLUT1^mEos2^ knock-in mice provides the unique opportunity to monitor precisely the relationship between VGLUT1 expression level and the quantal size of SV release. We thus, systematically investigated neurotransmission across various genotypes of VGLUT1^mEos2^ KI and VGLUT1 KO animals that express different levels of the VGLUT1 protein: WT (100%), VGLUT1^Eo/+^ (∼65%), VGLUT1^+/−^ (50%), VGLUT1^Eo/Eo^ (25%), and VGLUT1^−/−^ (0%).

First, we set to determine the contribution of VGLUT2 to excitatory synaptic transmission in our system. VGLUT2 is the primary isoform expressed in the VGLUT1 territory during pre-natal developmental stages (Miyazaki et al., 2003), and previous work has shown that a residual excitatory signal in VGLUT1−/− cells could be attributed to VGLUT2 expression in DIV10 to 20 neurons (Fremeau et al., 2004; Wojcik et al., 2004; Herzog et al., 2006). During post-natal development, VGLUT1 progressively overtakes VGLUT2 expression and reaches a plateau around the third post-natal week. In this study, to ensure that any observed effects are attributed solely to VGLUT1 expression changes and are not confounded by a compensatory role of VGLUT2, we investigated the temporal expression patterns of both VGLUT1 and VGLUT2 proteins in vitro. In dissociated cultured neurons from day in vitro (DIV) 12 through DIV 22 (Fig. 4A), we documented an increase in VGLUT1 expression concurrently with a decline in VGLUT2 levels which confirms our previous observation in immunocytochemistry (Zhang et al., 2019). By DIV 22, VGLUT2 expression was extremely low (Fig. 4A). we conclude that the switch between VGLUT1 and - 2 follows a similar spatiotemporal dynamic in cultures and tissue (Miyazaki et al., 2003; Wojcik et al., 2004; Zhang et al., 2019).

**Figure 4.**
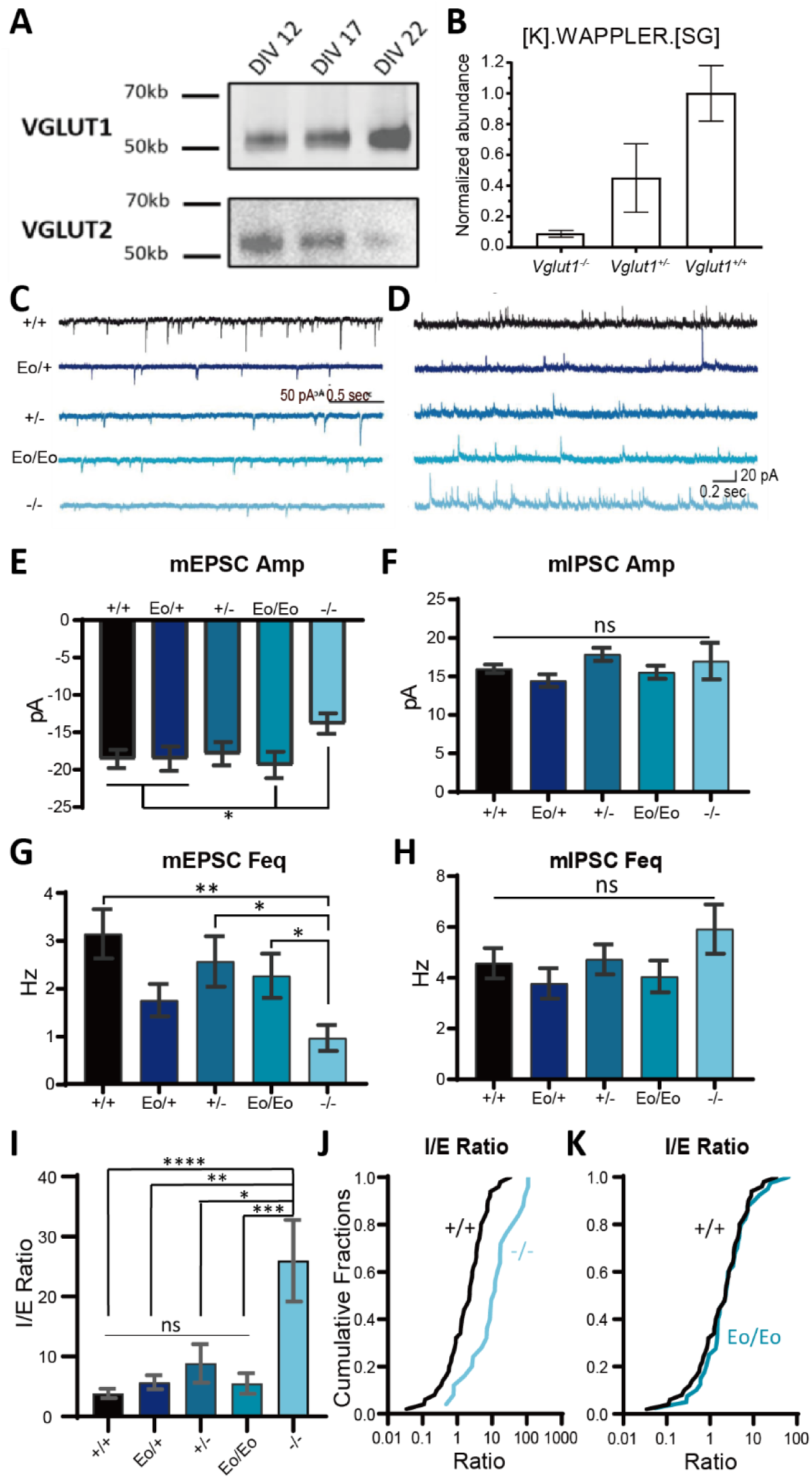
Decreasing levels of VGLUT1 have a minor impact on miniature neurotransmission compared to WT levels. (A) Western-blot assay reveals the progressive reduction of VGLUT2 expression across time (DIV) in hippocampal cultures. (B) MS/MS analysis show the peptide [K].WAPPLER.[SG] expression level in *Vglut1^−/−^*, *Vglut1^+/−^*, and *Vglut1^+/+^* hippocampal samples at P20. This peptide is common to VGLUT1 and −2. (C & D) Example traces of mEPSC and mIPSC recorded from the neuronal culture of different genotypes (WT: n = 50 cells, *Vglut1^Eo/+^*: n = 34 cells, *Vglut1^+/−^*: n = 35 cells, *Vglut1^Eo/Eo^*: n = 41 cells, and *Vglut1^−/−^*: n = 25 cells) that contain decreasing amounts of VGLUT1 from WT to null. (E & F) The amplitude of mEPSC and mIPSC. (G & H) The frequency of mEPSC and mIPSC. (I) The mIPSC/mEPSC frequency ratio of each genotype (J) Cumulative fractions of mIPSC/mEPSC frequency ratio of WT and *Vglut1^−/−^* cells. (K) Cumulative fractions of mIPSC/mEPSC frequency ratio of WT and *Vglut1^Eo/Eo^* cells. For (B), (E – H), (I), data are represented as mean ± SEM, * for *p* < 0.05, ** for *p* < 0.01, *** for *p* < 0.001, **** for *p* < 0.0001, ns for *p* > 0.05.

Next, we quantified the levels of VGLUT1 and VGLUT2 in mouse brains. We employed label free peptide quantification in mass spectrometry-based proteomics analysis, and evaluated the protein expression levels of VGLUT1 and VGLUT2 within the hippocampus of the different VGLUT1 KO genotypes at P20 (Fig. 4B, Table S1 – S3). Through numerous detections of isoform-specific peptides in VGLUT1 or VGLUT2, we were able to confirm that VGLUT2 expression remains unchanged, while VGLUT1 is reduced by 50% and 100% respectively in the VGLUT1 +/− and −/− animals (Table S1-S3). One peptide, [K].WAPPLER.[SG], that is shared by both VGLUT1 and −2, was detected at approximately 10% in VGLUT1^−/−^ compared to the expression level in VGLUT1^+/+^(Fig. 4B, Table S3). This residual presence suggests that it entirely represents VGLUT2, given the complete absence of VGLUT1 in these mice. In contrast, VGLUT1^+/−^ manifested a reduction to half the amount of this shared peptide compared to their +/+ counterparts (Fig. 4B). This strongly supports the notion that VGLUT1 is the predominant isoform within the hippocampus, with VGLUT2 expression represents roughly 10% of that of VGLUT1 in neuronal cultures after DIV20.

Next, we conducted an electrophysiological analysis in dissociated hippocampal cultures at DIV 17-21. Neurons from all genotypes had similar passive electrical properties (Fig. S2). We analyzed miniature excitatory postsynaptic currents (mEPSCs) and miniature inhibitory postsynaptic currents (mIPSCs; Fig. 4C, D), which serve as indicators of synapse maturation (Banerjee et al., 2021; Kavalali, 2015; Sutton et al., 2006). Strikingly, no significant differences in mEPSC amplitude were documented between neurons with varying degrees of VGLUT1 expression levels, except for VGLUT1^−/−^ full knock out neurons, where a significantly lower mEPSC amplitudes were recorded compared to WT neurons (Fig. 4E and S3). Importantly, even neurons with only 25% VGLUT1 expression (VGLUT1^Eo/Eo^) displayed mEPSC amplitudes equivalent to WT neurons.

Similarly, neurons with a full VGLUT1 knock out exhibited an appreciably lower frequency of mEPSCs in comparison to WT neurons, while in neurons with a progressive VGLUT1 knock down mEPSC frequencies were WT-like (Fig. 4G). As anticipated, neither the average mIPSC amplitude nor frequency exhibited significant differences across all genotypes (Fig. 4F, H), indicating no impairment of inhibitory synapses within these mutant lines. The residual mEPSC activity detected in VGLUT1^−/−^ neurons likely results from the remaining VGLUT2 expression (Fig. 4AB and Table S2).

Subsequently, we estimated the inhibitory-to-excitatory (I/E) balance within the networks from different genotypes to assess the relative contributions of excitation and inhibition (Fig. 4K). As expected, VGLUT1^−/−^ neurons, which exhibited a substantial reduction in the frequencies and amplitudes of mEPSCs (Fig. 4CD), had a significantly elevated I/E ratio compared to +/+ cultures (Fig. 4IJ). In contrast, the I/E ratios from VGLUT1 knock down cultures did not show any significant deviation from WT values (Fig. 4IK). We plotted the cumulative distribution of miniature events and I/E ratios for the two genotypes expressing the lowest levels of VGLUT1^−/−^ and VGLUT1^Eo/Eo^ *vs.* VGLUT1^+/+^ (Fig. S4; Fig. 4JK). The cumulative distribution curve of mEPSCs for VGLUT1^−/−^ neurons was notably shifted to the right compared to that of +/+ cells, indicating an overall decrease in excitatory synaptic activity (Fig. 4J). However, the I/E ratio curve for VGLUT1^Eo/Eo^ neurons, which retained only 25% of VGLUT1 expression, was barely distinguishable from that of WT neurons. This finding suggests that even a significantly reduced level of VGLUT1 is sufficient to maintain a relatively normal I/E balance (Fig. 4K). Collectively, these results demonstrate that excitatory quantal size is not linearly correlated to the expression level of the vesicular glutamate transporters and that only a full knock out of VGLUT1 shifts the I/E balance of dissociated neuronal cultures.

## Discussion

Here, we created and characterized a novel VGLUT1 knock-in mouse model tagged with the photoconvertible fluorescent reporter mEos2. Our aim was to develop a tool for tracking synaptic vesicle movements at an unprecedented level of detail using pointillist super-resolution techniques (sptPALM) as well as expanded distance and duration using mass photoconversion techniques. The VGLUT1^mEos2^ mouse line maintains normal VGLUT1 function and localization but shows reduced VGLUT1 expression levels. We took advantage of this adverse knock down phenotype to address a long-standing question regarding the relationship between VGLUT levels and quantal size modulation. Our data advocate for a non-linear relationship between VGLUT expression level and quantal size.

### VGLUT1^mEos2^ induces a knock-down of VGLUT1 but remains a suitable model for the imaging of synaptic vesicles dynamics

Traditionally, fluorescent reporters on SV membranes are easily introduced to neurons by transfection or viral transduction. However, concerns have been raised regarding potential overexpression artifacts. For example, a recent study showed that even moderate overexpression of SV protein will spill over proteins into dendrites and plasma membrane, which could lead to artefacts when examining protein localization (Watson et al., 2023). Thus, knock-in strategies provide the advantage to potentially avoid such problems.

Indeed, VGLUT1^mEos2^ was correctly targeted to synaptic sub-compartment throughout the brain and displayed a punctate localization at thin neurites like VGLUT1. We provide some evidence that VGLUT1^mEos2^ is a functional transporter as previously shown for VGLUT1^venus^. Surprisingly though, tagging VGLUT1 with mEos2 led to a significant decrease in VGLUT1 expression. This is despite the fact that we employed the same strategy used in creating the VGLUT1^venus^ KI mice (Herzog et al., 2011), where no expression level changes were detected. By monitoring the ratio between VGLUT1^mEos2^ and WT during postnatal development via qPCR, we show that reduction in VGLUT1 mRNA levels are the cause for lower VGLUT1 expression. Similarly, tagging PSD-95 with mEos2 resulted in knockdown of PSD-95 in the PSD-95^mEos2^ KI mouse line (Broadhead et al., 2016). In contrast, Venus-tagged VGLUT1 and Munc18-1 showed no such effect (Cijsouw et al., 2014; Herzog et al., 2011). While both mEos2 and Venus are optimized for mammalian expression, Venus is an improved GFP variant designed for rapid synthesis, folding, and low toxicity, whereas mEos2 is a first-generation EosFP derivative optimized for reduced acid sensitivity and photostability (McKinney et al., 2009; Nagai et al., 2002). In recent years, several variants derived from mEos2 have emerged through distinct point mutations, such as mEos3.2 and mEos4a/b (Paez-Segala et al., 2015; Zhang et al., 2012). Although these new fluorescent proteins have been extensively characterized in terms of their photo-physical properties (Berardozzi et al., 2016; Bourgeois, 2023; Thédié et al., 2017; Wulffele et al., 2022), it is yet to be determined if they exert similar consequences to protein synthesis. Notably, no existing research has satisfactorily explained the reason behind the suppressive effect of mEos2 for mRNA expression. To bridge this knowledge gap, mRNA transcription and degradation mechanisms must be explored, which falls beyond the scope of this study.

The established cellular and molecular characterization of VGLUT1^+/−^ mice revealed very mild impact of VGLUT1 haploinsufficiency on neurotransmission and morphology (Fremeau et al., 2004; Wojcik et al., 2004). Hence, VGLUT1^Eo/+^ with a moderate knockdown effect above 60% expression level compared to WT seem a good compromise to study SV super-pool in near-native conditions. Furthermore, the fluorescent signal in VGLUT1^Eo/+^ is sufficient to monitor SV mobility under various paradigms.

### Tracing axonal synaptic vesicle mobility using VGLUT1^mEos2^

At the presynaptic terminal, various SV pools displaying different release propensities have been characterized (Sudhof, 2004; Rizzoli and Betz, 2005). However, the characterization of the localization and mobility properties of these different SVs is still lacking high spatial resolution. Combined with electron microscopy techniques, releasable SVs were revealed to have extremely intermixed localizations inside the boutons (Rizzoli and Betz, 2004; Darcy et al., 2006)and ultrafast endocytosis through intermediate endosomes (Watanabe et al., 2014; Watanabe & Boucrot, 2017). Additionally, beyond presynaptic terminals, SVs exchange through a super-pool spanning several *en-passant* boutons of the axon (Darcy et al., 2006; Staras et al., 2010; Herzog et al., 2011; Zhang et al., 2019).

We demonstrate here that sptPALM and mass photoconversion experiments enable the investigation of SV dynamics in the axon. The sptPALM technique allowed to track individual VGLUT1 molecules and thus single SVs with unprecedented precision. We foresee that it will enable to dissect distinct stages of SV mobility in the axon and terminals. This includes the monitoring of pure axonal transport paths, extra-synaptic movements, and the critical transitions between axonal transport and synaptic clustering, as well as the reverse transition from synaptic clusters back to axonal transport. Conversely, with mass photoconversion experiments, we gain the capacity to observe neuron-wide or axon-specific SV exchanges over extended spatio-temporal scales. Here we could monitor the evolution of 5 boutons of the same axon over 6 hours and 40 minutes with the repeated photoconversion at another bouton of this same axon. Photoconverted VGLUT1^mEos2^, ended up populating all boutons monitored. Even when a SV cluster appeared to be diminishing (Fig S1 boutons 1 and 2), there persisted a continuous process of integration of photoconverted SVs. In the future the VGLUT1^mEos2^ knock-in will allow a wide range of experimental paradigms both in-vivo and in-vitro.

### VGLUT1 expression level has a minor impact on quantal size

Experimental evidence has consistently demonstrated for several neurotransmitters that modulating vesicular transporter levels significantly impacts the quantal size (Song et al., 1997; Caudle et al., 2007; Lohr et al., 2014; Taylor et al., 2014). For excitatory synapses, the situation seems slightly more complex. In drosophila, dVGLUT over-expression led to an increased quantal size (Daniels et al., 2004), and the knock-down of VGLUT2 in mice leads to reduced mEPSC amplitudes in thalamic neurons (Moechars et al., 2006). However, while overexpression of VGLUT1 in mouse models lead to increased amplitudes of mEPSCs, heterozygous deletion of VGLUT1 had no effect on mEPSC amplitudes compared to WT littermate cultures (Weston et al., 2011; Wojcik et al., 2004). In VGLUT1 KO neurons, the average mEPSCs amplitude is smaller and the mEPSC frequency is lower, with the residual activity attributed to VGLUT2 expression in hippocampal neurons (Wojcik et al., 2004). Conversely, some reports suggest only reduced mEPSC frequency but no change in the amplitude of the remaining VGLUT2 events (Fremeau et al., 2004). The present data provide a fresh and thorough survey to these conflicting data. We implemented a precise quantification of VGLUT2 expression in the hippocampus during the third post-natal week. We conclude that VGLUT2 represents around 10% of VGLUT1 at this age in vivo and in vitro. To date, the best estimates for the number of VGLUT molecules are between 5 and 15 molecules per SV (Mutch et al., 2011; Takamori et al., 2006; Wilhelm et al., 2014), and we have previously shown that VGLUT1 and −2 can reside on the same vesicles in the hippocampus (Herzog et al., 2006). Based on this, we estimate that VGLUT1^−/−^ cells should have between 0 and 1 VGLUT2 transporter per SV. Our experiment tested four conditions of VGLUT1 expression (100%, 65%, 50%, 25% and 0%). The absence of reduction in the quantal size in all knock-down conditions, and that only a 25% reduction in the mEPSC amplitude was documented in the full knock-out, advocates for a very mild impact of VGLUT1 level on quantal size in hippocampal neurons. These data also fit with the model proposed in drosophila neuromuscular junction suggesting that a single dVGLUT molecule is sufficient to load SVs with adequate glutamate (Daniels et al., 2006). The reduction in mEPSC frequency is also in agreement with the likely occurrence of many SVs baring no VGLUT on their membrane (Fremeau et al., 2004; Wojcik et al., 2004; Daniels et al., 2006). Yet the difference of relationship between VGLUT and quantal size observed between hippocampal neurons expressing mostly VGLUT1 and thalamic neurons expressing mostly VGLUT2 is intriguing and calls for a more thorough survey of this comparison between the two transporters and the two neuronal populations (Fremeau et al., 2004; Wojcik et al., 2004; Moechars et al., 2006). While the plateau of loading might be poorly correlated to the VGLUT level, investigations at the calyx of Held synapses that express both VGLUT1 and −2 revealed that VGLUT1 loss slows down SV refilling with glutamate to an extent that can be rate limiting for neurotransmission (Nakakubo et al., 2020).

VGLUT1^−/−^ mice begin to exhibit a phenotype in the third postnatal week, as VGLUT2 levels decline. This phenotype becomes severe within days, and they die around 18 days after birth (Wojcik et al., 2004). Studies using VGLUT1^+/−^ mice show mild phenotypes in spatial learning tasks, a deficit in long-term potentiation (LTP) in the CA1 region (Balschun et al., 2010) and anxiety/depression related behaviors (Garcia-Garcia et al., 2009). The latter was attenuated after a chronic treatment with conventional antidepressants but not with ketamine (Belloch et al., 2023). The homozygote of VGLUT1^mEos2^ doesn’t show any obvious behavioral changes and are healthy. We therefore conclude that the residual expression of VGLUT1 to 25% of WT levels are sufficient for mouse survival, but a more detailed behavioral analysis will be needed to consolidate a relationship between VGLUT1 expression levels and mouse behavior. We anticipate that VGLUT1^Eo/Eo^ will provide deeper insights into how VGLUT1 deficits contribute to an imbalance between glutamate and monoamines signalling that translates into pathological behavioral phenotypes.

Overall, the VGLUT1^mEos2^ mice constitute an unexpected knockdown model of the expression of VGLUT1. Yet we show that it remains a promising tool for the investigation of both synaptic vesicle mobility and the impact of hypo-glutamate function on brain physiology. We took advantage of this new model to reveal that VGLUT1 level at hippocampal synapses is not a major regulator of quantal size, which may not be a general rule at excitatory synapses. The mechanisms behind these observations remain to be unraveled.

## Methods

### Animals

All mice are housed in 12/12 LD with ad libitum feeding. Every effort was made to minimize the number of animals used and their suffering. The experimental design and all procedures were in accordance with the European guide for the care and use of laboratory animals and approved by the ethics committee of Bordeaux Universities (CE50) under the APAFIS n°1692.

### Antibodies

The primary antibodies used were: VGLUT1 rabbit polyclonal antiserum BN3L2Bf (Herzog et al., 2001), Snap 25 mouse monoclonal antibody (111011, SYSY). The secondary antibodies were: Donkey anti-rabbit Alexa 488 (A21206, Invitrogen), goat anti-mouse Alexa 488 (A32723, Invitrogen) and goat anti-rabbit IRDye 800CW (P/N 925-32211, Li-COR).

### Generation of VGLUT1^mEOS2^ KI mice

The targeting vector was constructed on the basis of a 14 kb genomic clone of the *VGLUT1* locus in pBluescript, which had been isolated from a λFIXII genomic library of the SV129 mouse strain (Stratagene). In the targeting vector, the STOP codon in the last exon (exon 12) of the *VGLUT1* gene was replaced in-frame by a *mEOS2-loxP-neo-loxP* cassette using recombineering techniques with engineered primers. The *venus-loxP-neo-loxP* cassette was inserted between a 6.7 kb genomic sequence in 5’ position and a 7.9 kb genomic sequence in 3’ position. Mice carrying the mutant *VGLUT^mEOS2-neo^*gene (*VGLUT1^Eon/+^*) were generated by homologous recombination in embryonic stem cells (SV129/ola) and identified by Southern blotting. To eliminate deleterious effects of the neomycin resistance gene, we crossed heterozygous *VGLUT1^Eon/+^* mice with Ella-Cre mice that express Cre recombinase in early embryonic stages (Lakso et al., 1996). Successfully recombined *VGLUT1^mEos2^* alleles (Eo/+) in offspring from these interbreedings were genotyped by PCR. *Slc17a*7 gene is located on the mouse chromosome 7 contig NT_039424.7 at position 6081938-6082223. The following oligonucleotides were used for genotyping PCRs: 9420, CTGGCTGGCAGTGACGAAAG; 9421, CGCTCAGGCTAGAGGTGTATGGA; 32896, CTGAAGTCACATCGGTAATG. Oligonucleotides 9420 and 9421 were used to amplify the *VGLUT1^+^* and *VGLUT1^Eo^* allele, with expected band size 202 bp and 1033 bp respectively. Oligonucleotides 9420 and 32896 were used to amplify the *VGLUT1^Eo^* allele with an expected band of 668 bp.

### Hippocampal cell culture

Hippocampal primary dissociated cultures were prepared from P0 pups from different genotypes. The hippocampi were dissected in ice-cold Leibovitz’s L-15 medium (11415064; Gibco), and then incubate in 0.05% trypsin-EDTA (25300054, Gibco) for 15 min at 37°C. The tissues were washed with Dulbecco’s Modified Eagle’s Medium (DMEM, 61965026, Gibco) containing 10% FBS (CVFSVF0001, Eurobio), 1% Penicillin-streptomycin (15140122, Gibco). Cells were mechanically dissociated by pipetting up and down. Cells were plated onto poly-L-lysine (P2636, Sigma) coated coverslips at a density of 20 000 cells/cm^2^. Cells were grown in Neurobasal A medium (12349105, Gibco) containing 2% B27 supplement (17504044, Gibco), 0.5 mM Glutamax(35050038, Gibco), and 0.2% MycoZap plus-PR (VZA2021, Lonza). After 10 days, half of the medium was changed. Imaging and electrophysiological recording of live dissociated neuron cultures was performed at 17-21 days *in vitro* (DIV).

In the PALM imaging experiments, neurons were transduced at DIV1 with Lenti virus of F(syn)W-RBN::VGLUT1mEos2.

In the massive photoconversion experiments, the VGLUT1^Eo/+^ and VGLUT1^+/+^ cells were plated with a ratio of 1:10 to better isolate the fluorescent synapse with VGLUT1^mEos2^.

### Biochemical analyses

Sodium dodecyl sulfate polyacrylamide gel electrophoresis (SDS-PAGE) and Western blotting were performed according to standard procedures using Alexa-488 or IRDye 800-coupled secondary antibodies for semiquantitative Western detection. Alexa 488 secondary antibodies were used for the VGLUT1 detection in different VGLUT1^mEos2^ genotypes and fluorescent signals were visualized with ChemiDoc MP system (Bio-Rad). Western blotting for VGLUT2 served as a loading control for VGLUT1 and VGLUT^mEos2^. The IRDye 800 secondary antibody was used for the developmental pattern of VGLUT1^mEos2^ and signals were visualized with Infrared Odyssey (Li-COR Biosciences). Quantitative data are expressed as mean ± SEM.

### Imaging of VGLUT1^mEos2^ brain

Direct VGLUT1^mEos2^ fluorescence was performed on perfusion-fixed (4% paraformaldehyde) brain or 60 µm brain section. Pictures were taken with epi-fluorescent microscope (Nikon, ND2). The local photoconversion was done with the DAPI filter under a 40× objective. The duration of UV light exposure was 10 seconds at full intensity of the Intensilight source.

### PALM imaging

For PALM imaging, the acquisitions were performed on an inverted motorized microscope Nikon Ti Eclipse (Nikon France S.A.S., Champigny-sur-Marne, France) equipped with a 100× 1.49 NA PL-APO objective with a perfect focus system for axial stabilization. The fluorescence signal was collected by the objective and focused onto a sensitive Evolve EMCCD camera (Photometrics, Tucson, USA) through the combination of a dichroic and emission filters (D101-R561 and F39–617, respectively, Chroma). The entire setup was packaged in a temperature-controlled chamber setting to 37°C (Life Imaging Services, Basel, Switzerland). VGLUT1^mEos2^ was photoactivated using a 405 nm laser (3 W/cm^2^), and the resulting photoconverted single molecule fluorescence was excited with a 561 nm laser (0.8 kW/cm^2^). Both lasers illuminated the sample simultaneously. The acquisition was driven by MetaMorph (Molecular Devices, Sunnyvale, USA) in streaming mode at 20 frames per second (50 ms exposure time) for 3,000 frames, each 256×256 pixels. These image stacks were then processed offline for single molecule localization and tracking using the PALMTracer software, a MetaMorph (Molecular Devices, Sunnyvale, USA) add-on developed at the Interdisciplinary Institute of Neuroscience (IINS -UMR5297 -CNRS / University of Bordeaux), by Corey Butler (IINS -UMR5297 -CNRS / University of Bordeaux), Adel Mohamed Kechkar (Ecole Nationale Supérieure de Biotechnologie, Constantine, Algeria) and Jean-Baptiste Sibarita (IINS -UMR5297 -CNRS / University of Bordeaux).

Briefly, Single molecule localization was achieved using wavelet segmentation, and then filtered out based on the quality of a 2D Gaussian fit. SPT analysis was then done based on the detections using reconnection algorithms and for MSD and D calculations on reconnected trajectories (Izeddin et al., 2012; Kechkar et al., 2013; Racine et al., 2007). Super resolution tracking images were reconstructed here with a pixel size of 40 nm.

### Mass photoconversion at individual boutons

Photoconversion was performed with the neuronal culture to check the SVs trafficking along the axons. The imaging acquisition was done with a spinning-disk confocal microscope that was a Leica DMI6000 with an autofocus system (Leica Microsystems, Wetzlar, Germany), equipped with a confocal Scanner Unit CSU-X1 (Yokogawa Electric Corporation, Tokyo, Japan), and an Evolve EMCCD camera (Photometrics, Tucson, USA). The photoconversion experiments were done with a FRAP scanner system (Roper Scientific, Evry, France). Surrounding the setup, a thermal incubator is set to 37°C (Life Imaging Services, Basel, Switzerland).

Several presynaptic boutons with VGLUT1^mEos2^ molecules on isolated axons were randomly selected to photoconvert. The 488 nm laser (15 mW) and 561 nm laser (15 mW) were used to monitor the fluorescence of VGLUT1^mEos2^ emitting green or red light respectively. To obtain enough converted mEos2 molecules, the photoconversion was achieved though 40 passes of 405 nm laser (80 mW).

Image acquisition was monitored every 2 s for 4 s before the photoconversion, and every 5 min after the photoconversion during the next 40 min. Then the photoconversion cycle was repeated 9 times. To photoactivate dark mEos2 molecules, a 300 ms exposure with 405 nm (10 mW) was applied at the beginning of the acquisition. This had no measurable impact on fluorescence levels in the red channel. The entire procedure was controlled by MetaMorph software (Molecular Devices, Sunnyvale, USA).

### Proteomics

#### Sample preparation and protein digestion

Proteomics were performed as previous reported (Blumenstock et al., 2019). Protein samples were solubilized in Laemmli buffer and separated using SDS-PAGE. After colloidal blue staining, the gel weight range between 50-70 kD was collected and subsequently cut in 1 mm x 1 mm gel pieces. Gel slices were destained in 25 mM ammonium bicarbonate 50% acetonitrile (ACN), shrunk in ACN for 10 min. After ACN removal, gel pieces were dried at room temperature, covered with trypsin solution (10 ng/µl in 40 mM NH_4_HCO_3_ and 10% ACN), rehydrated at 4 °C for 10 min and incubated overnight at 37 °C. Spots were then incubated for 15 min in 40 mM NH_4_HCO_3_ and 10% ACN at room temperature with rotary shaking. The supernatant was collected and an H_2_O/ACN/HCOOH (47.5:47.5:5) extraction solution was added onto gel pieces for 15 min. The extraction step was repeated twice. Supernatants were pooled and concentrated in a vacuum centrifuge before being resuspended in 30 µl 0.1% formic acid.

#### nLC-MS/MS analysis and Label-Free Quantitative Data Analysis

Peptide mixture was analyzed on a Ultimate 3000 nanoLC system (Dionex, Amsterdam, The Netherlands) coupled to a Electrospray Orbitrap Fusion™ Lumos™ Tribrid™ Mass Spectrometer (Thermo Fisher Scientific, San Jose, CA). Ten microliters of peptide digests were loaded onto a 300-µm-inner diameter x 5-mm C_18_ PepMap^TM^ trap column (LC Packings) at a flow rate of 10 µL/min. The peptides were eluted from the trap column onto an analytical 75-mm id x 50-cm C18 Pep-Map column (LC Packings) with a 5–27.5% linear gradient of solvent B in 105 min (solvent A was 0.1% formic acid and solvent B was 0.1% formic acid in 80% ACN) followed by a 10 min gradient from 27.5% to 40% solvent B. The separation flow rate was set at 300 nL/min. The mass spectrometer operated in positive ion mode at a 2-kV needle voltage. Data were acquired using Xcalibur 4.3 software in a data-dependent mode. MS scans (*m/z* 375-1500) were recorded in the Orbitrap at a resolution of R = 120 000 (@ m/z 200) and an AGC target of 4 × 10^5^ ions collected within 50 ms. Dynamic exclusion was set to 60 s and top speed fragmentation in HCD mode was performed over a 3 s cycle. MS/MS scans were collected in the Ion trap with a maximum fill time of 300 ms. Only +2 to +7 charged ions were selected for fragmentation. Other settings were as follows: no sheath nor auxiliary gas flow, heated capillary temperature, 275 °C; normalized HCD collision energy of 30%, isolation width of 1.6 m/z and AGC target of 3 × 10^3^. Monoisotopic precursor selection (MIPS) was set to Peptide and an intensity threshold was set to 5 × 10^3^.

#### Database search and results processing

Data were searched by SEQUEST through Proteome Discoverer 2.2 (Thermo Fisher Scientific Inc.) against a *Mus musculus* uniprot database (53,378 entries in v2018-08). Spectra from peptides higher than 5000 Da or lower than 350 Da were rejected. Search parameters were as follows: mass accuracy of the monoisotopic peptide precursor and peptide fragments was set to 10 ppm and 0.6 Da respectively. Only b- and y-ions were considered for mass calculation. Sequest HT was used as the search algorithm : Oxidation of methionines (+16 Da) and protein N-terminal acetylation (+42 Da) were considered as variable modifications while carbamidomethylation of cysteines (+57 Da) was considered as fixed modification. Two missed trypsin cleavages were allowed. Peptide validation was performed using Percolator algorithm (Käll et al., 2007) and only “high confidence” peptides were retained corresponding to a 1% False Positive Rate at peptide level. Peaks were detected and integrated using the Minora algorithm embedded in Proteome Discoverer. Normalization was performed based on total peptide amount. Protein ratio were calculated as the median of all possible pairwise peptide ratios. A t-test was calculated based on background population of peptides or proteins. Quantitative data were considered for proteins quantified by a minimum of two peptides and a statistical p-value lower than 0.05.

### Electrophysiology

Whole-cell patch-clamp recordings were performed from DIV17 to DIV21. Patch pipettes (2-4MΩ) were filled with the following intracellular solution (in mM): 125 CsMeSO_3_, 2 MgCl_2_, 1 CaCl_2_, 4 Na_2_ATP, 10 EGTA, 10 HEPES, 0.4 NaGTP and 5 QX-314-Cl, pH was adjusted to 7.3 with CsOH. Extracellular solution was a standard ACSF containing the following components (in mM): 124 NaCl, 1.25 NaH_2_PO_4_, 1.3 MgCl_2_, 2.7 KCL, 26 NaHCO_3_, 2 CaCl_2_, 18.6 Glucose and 2.25 Ascorbic acid. To record excitatory and inhibitory miniature currents (mEPSC and mIPSC), Tetrodotoxin (TTX) was added at 1µM into an aliquot of the standard ACSF (Alomone labs). Cultures were perfused at 35°C with an ACSF perfusion speed of 0.02mL/min and equilibrated with 95% O2/5% CO_2_. Signals were recorded at different membrane potentials under voltage clamp conditions for about 2 min (0mV for inhibitory events and −70mV for excitatory events) using a MultiClamp 700 B amplifier (Molecular Devices, Foster City, CA) and Clampfit software. Recording at 0 mV in voltage clamp allowed us to confirm the effect of TTX on network activity. The recording of miniature events began 2 min afterADDINg TTX and the extinction of synchronized IPSCs. Additional recordings were performed at membrane potentials of −20 mV, −40 mV, −60 mV and −80 mV. For drug treatment, CNQX was used at 50 µM and PTX at 100 µM and 2 minutes after drug addition, the condition was considered as stable.

### Electrophysiology Analysis

Analyses were performed using Clampfit (Molecular Device), in which we created one mini-excitatory (−70 mV) and one mini-inhibitory (0 mV) template from a representative recording. Those templates were used for all recordings and analysis was done blind to the experimental group. We measured the number of mEPSCs at −70 mV and mIPSCs at 0 mV and their mean amplitude. Cell properties were monitored to get a homogenous set of cells, i.e. we analyzed the seal-test recordings of every cell (Fig S3) and calculated the capacitance from the Tau measured by Clampfit (Fig S3). Cells with a leak current over −200pA, and/or a membrane resistance over −100MOhms were excluded from the analysis. Statistical analyses were performed using One-way ANOVA or Kruskal-Wallis (* for p < 0.05, ** for p < 0.01 and *** for p < 0.001).

## Acknowledgement

We express our deep gratitude to Nils Brose for his invaluable support in the production of the VGLUT1^mEos2^ KI line, his assistance with technical troubleshooting, and his insightful discussions throughout the project. We thank Dilja Krueger-Burg, Fritz Benseler, Astrid Zeuch, Christel Poujol, Elizabeth Normand, Pierre Costet, the AGCT Lab, the MPIEM animal facility for excellent technical support. Serge Marty and Martin Oheim for fruitful discussions. Sonja M. Wojcik for sharing the VGLUT1 knock-out mouse line. Several experiments required the use of Bordeaux University/CNRS/INSERM core facilities: Bordeaux Imaging Center (member of France BioImaging supported by the French National Research Agency; ANR-10-INBS-04); Vect’UB viral vector facility; Biochemistry and Biophysics of Proteins core facility (BioProt); Mouse breeding facility (SCA); Bordeaux Proteome. X.M.Z. was supported by the Erasmus Mundus ENC program, the National Natural Science Foundation of China (32100761), Guangdong Basic and Applied Basic Research Foundation (2024A1515013101), and Guangzhou Basic and Applied Basic Research Foundation (SL2024A04J01782). X.M.Z. and U.F. were supported by the labex BRAIN extension grant (ANR-10-LABX-43 BRAIN). Funding from the Agence Nationale de la Recherche (ANR-12-JSV4-0005-01 VGLUT-IQ ; ANR-10-LABX-43 BRAIN; ANR-10-IDEX-03-02 PEPS SV-PIT to E.H.). M.F.A. was supported by the Fondation pour la Recherche Médicale (ING20150532192 respectively). This work was supported by the German Research Foundation Excellence Strategy EXC-2049-390688087 (N.L); CRC 1286 “Quantitative Synaptology” project A11 (N. L).

## Author contributions

X.M.Z, performed experiments, analyzed results and wrote the manuscript; U.F., M.F.A, S.C, M.M., C.M., performed experiments, analyzed results; M.D. managed the animal breeding; Y.H. conceived experiments, analyzed results and edited the manuscript; NL: conceived and supervised experiments, analyzed results and edited the manuscript; E.H. supervised and funded the project, conceived experiments, analyzed results, wrote parts of the manuscript. All authors reviewed the final versions of the manuscript.

---

**Supplementary figure 1, related to figure 3.**
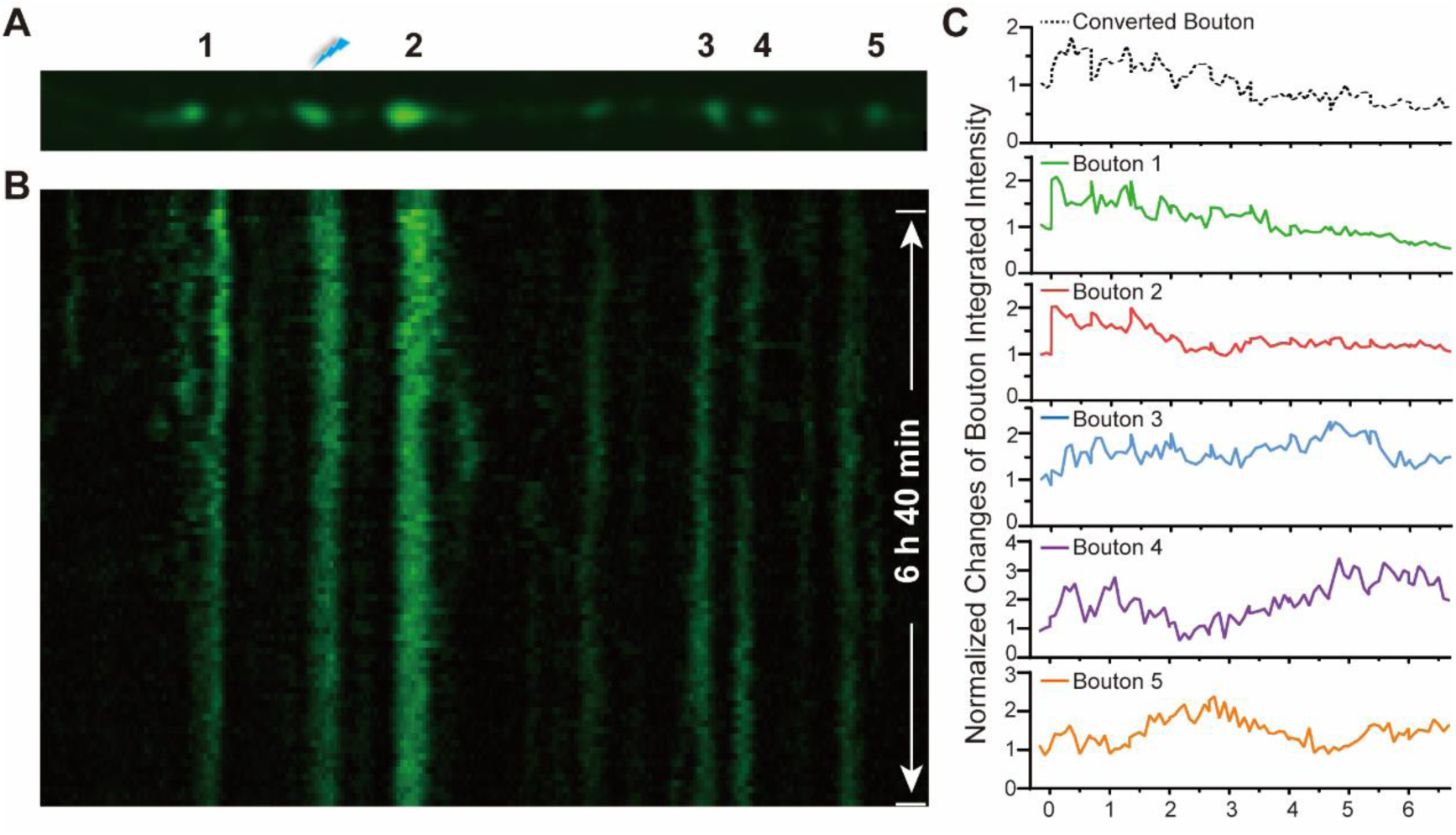
Green channel images of photoconversion at *en passant* boutons. (A) Imaging of an axonal fiber with 6 *en passant* boutons. One synapse was selected and photoconverted with 405 nm laser. (B) Kymographs of the live imaging acquisition in the green channel. (C) The intensity plot of each monitored synapse with 488 nm laser excitation.

**Supplementary figure 2.**
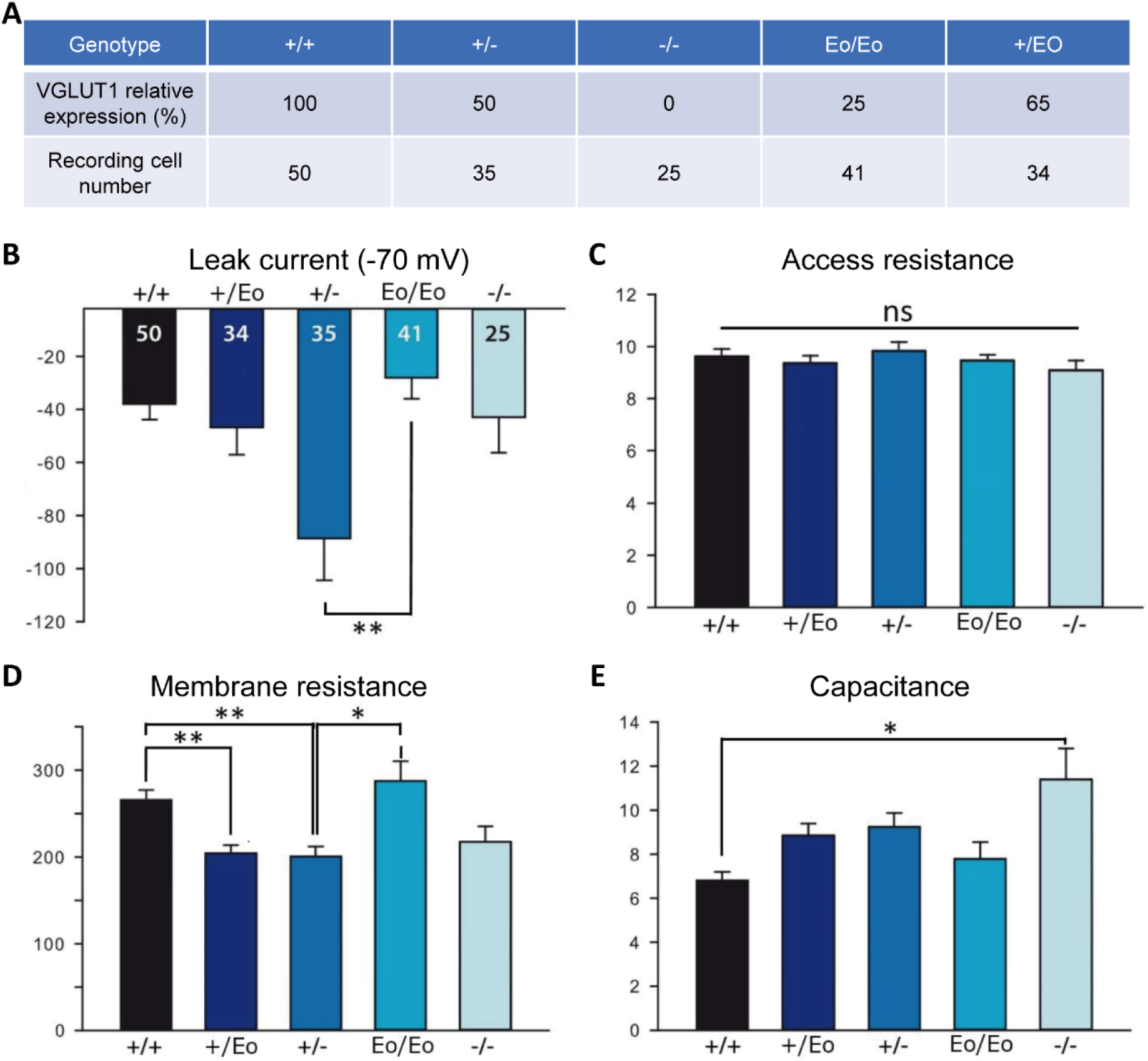
The properties of neurons examined in electrophysiology test. (A) The table shows the VGLUT1 relative expression level and the cell numbers recorded from different genotypes. (B-E) Seal test to measure the cell properties of the leak current (B), the access resistance (C), the membrane resistance (D), and the capacitance (E).

**Supplementary figure 3.**
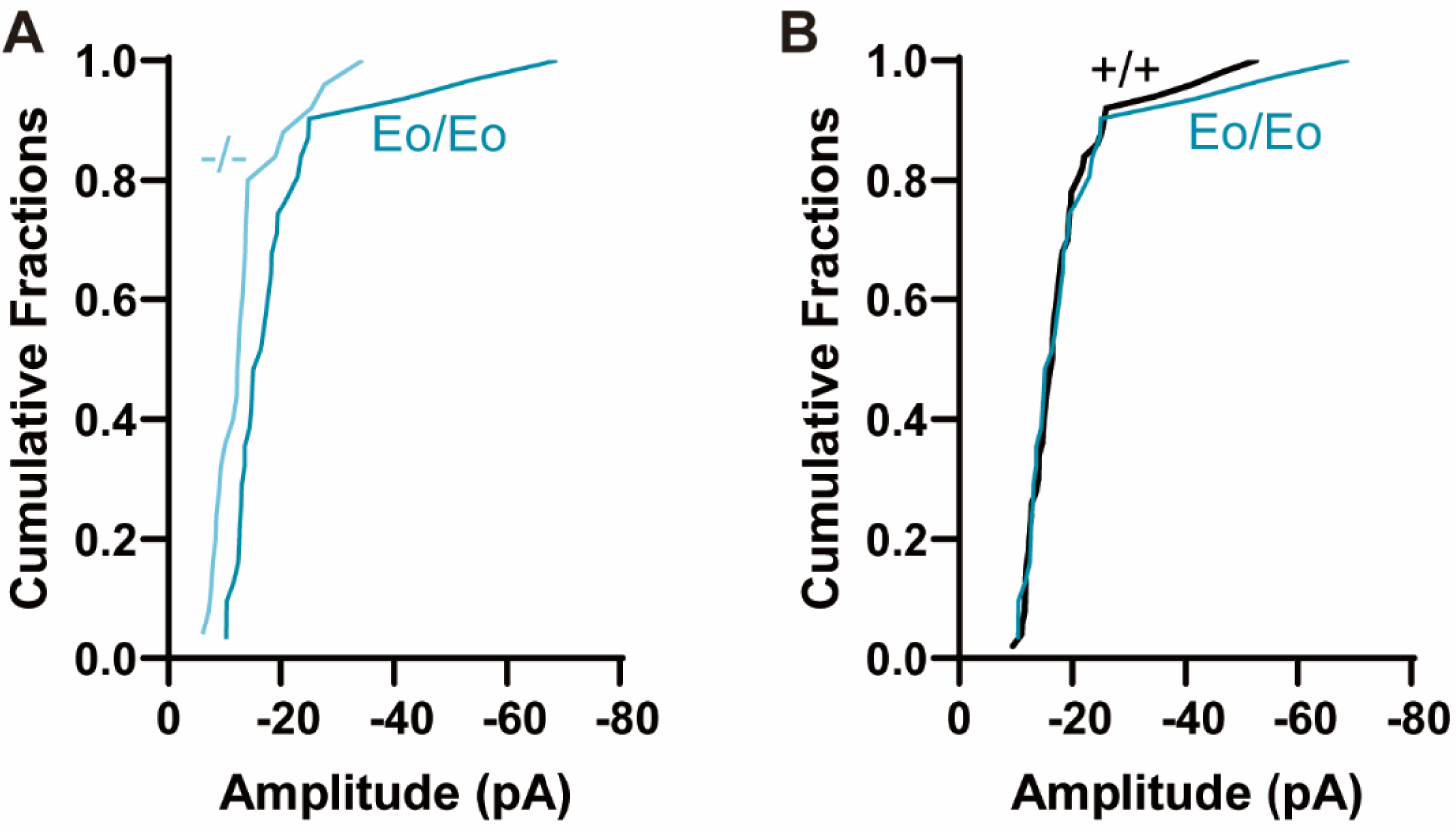
Severe knockdown of VGLUT1 does not impact the cumulative distribution of mEPSC amplitudes. The cumulative fraction of mEPSC frequency recorded from VGLUT1^−/−^ *vs.* VGLUT1^Eo/Eo^ neuronal culture (A), and VGLUT1^+/+^ vs. VGLUT1^Eo/Eo^ neuronal culture (B).

**Supplementary figure 4.**
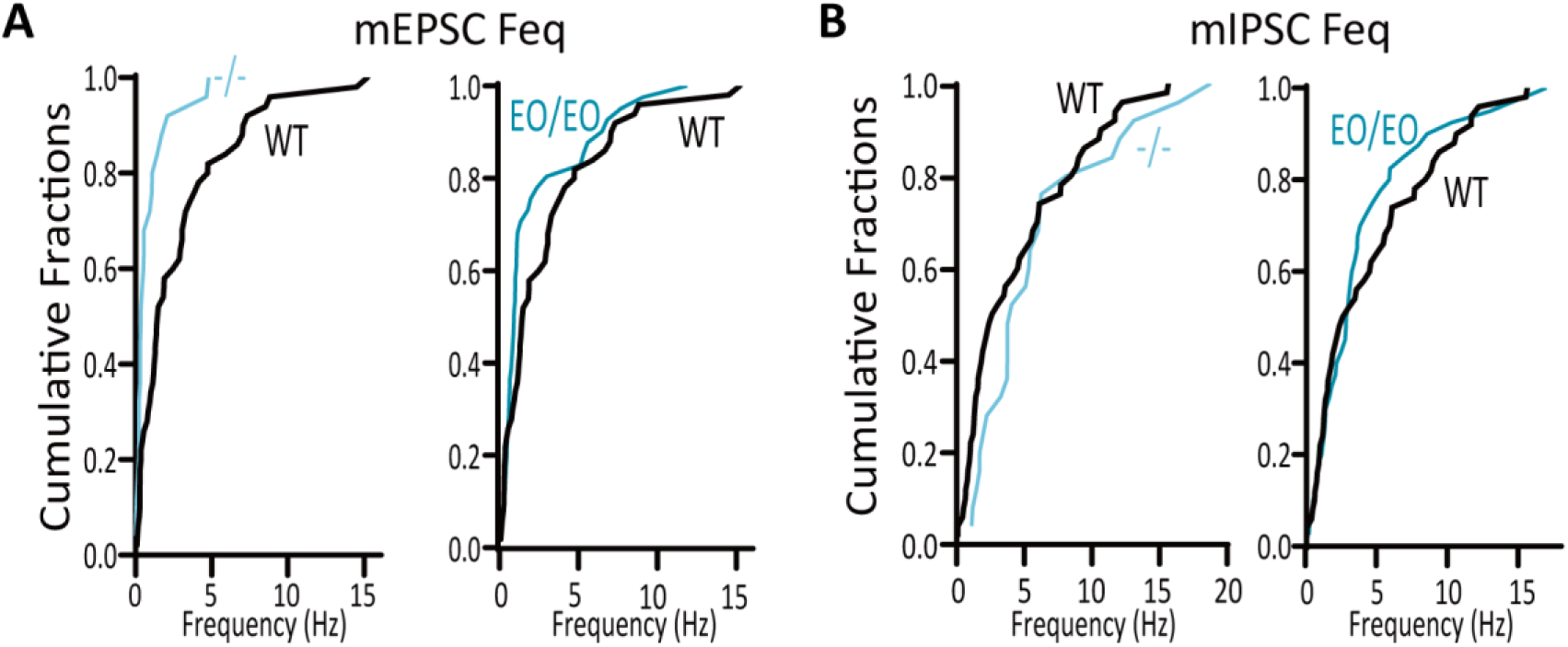
The absence of VGLUT1 expression impairs the mEPSC frequency, but not the mIPSC frequency. (A) The cumulative fraction of mEPSC frequency was recorded from WT, *Vglut1^Eo/Eo^*, and *Vglut1^−/−^*neuronal culture. (B) The cumulative fraction of mIPSC frequency was recorded from WT, *Vglut1^Eo/Eo^*, and *Vglut1^−/−^* neuronal culture.

**Table S1.**

| VGLUT1 specific peptide |  |  |
| --- | --- | --- |
| Peptide | Abundance Ratio |  |
|  | *Vglut1*<sup>+/-</sup> / *Vglut1*<sup>+/+</sup> | *Vglut1*<sup>-/-</sup> / *Vglut1*<sup>+/+</sup> |
| [R].DPPVVDCTCFGLPR.[R] | 0.202 | 0.001 |
| [K].QPWAEPEEMSEEK.[C] | 0.444 | 0.001 |
| [R].QEGAETLELSADGRPVTTHTR.[D] | 0.474 | 0.001 |
| [R].KYIEDAIGESAK.[L] | 0.288 | 0.001 |
| [K].RQEGAETLELSADGRPVTTHTR.[D] | 0.293 | 0.001 |
| [K].QPWAEPEEMSEEK.[C] | 0.319 | 0.001 |
| [R].HIMSTTNVR.[K]* | 0.495 | 0.001 |
| [R].HIMSTTNVR.[K] | 0.57 | 0.001 |
| [K].YIEDAIGESAK.[L] | 0.444 | 0.017 |
| [K].LMNPVTK.[F] | 0.432 | 0.001 |
| Mean±SEM | 0.396 ± 0.114 | 0.003 ± 0.005 |

Note: \* with 1×oxidation modification.

**Table S2.**
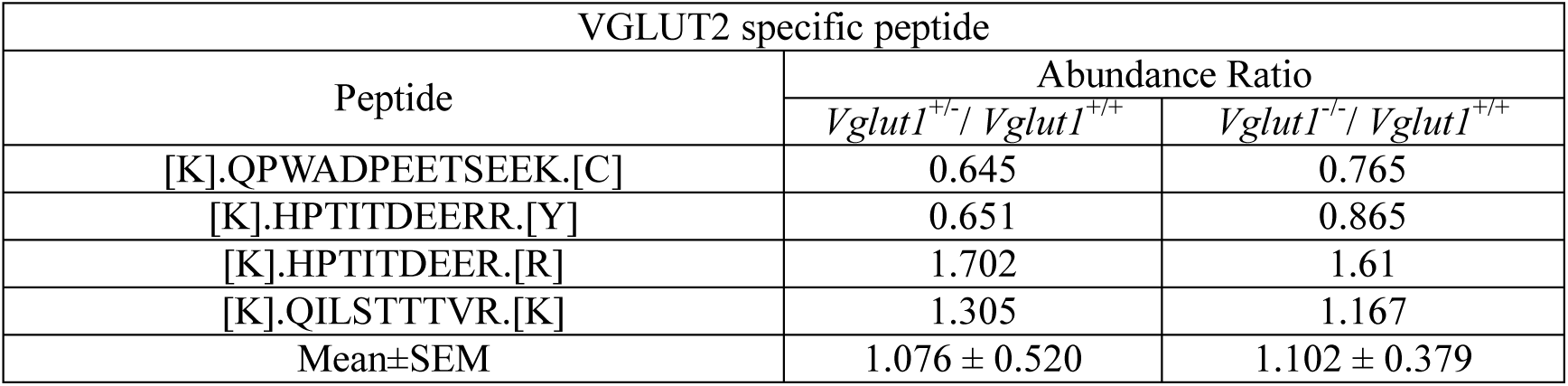

| VGLUT2 specific peptide |  |  |
| --- | --- | --- |
| Peptide | Abundance Ratio |  |
|  | *Vglut1*<sup>+/-</sup> / *Vglut1*<sup>+/+</sup> | *Vglut1*<sup>-/-</sup> / *Vglut1*<sup>+/+</sup> |
| [K].QPWADPEETSEEK.[C] | 0.645 | 0.765 |
| [K].HPTITDEERR.[Y] | 0.651 | 0.865 |
| [K].HPTITDEER.[R] | 1.702 | 1.61 |
| [K].QILSTTTVR.[K] | 1.305 | 1.167 |
| Mean±SEM | 1.076 ± 0.520 | 1.102 ± 0.379 |

**Table S3.**
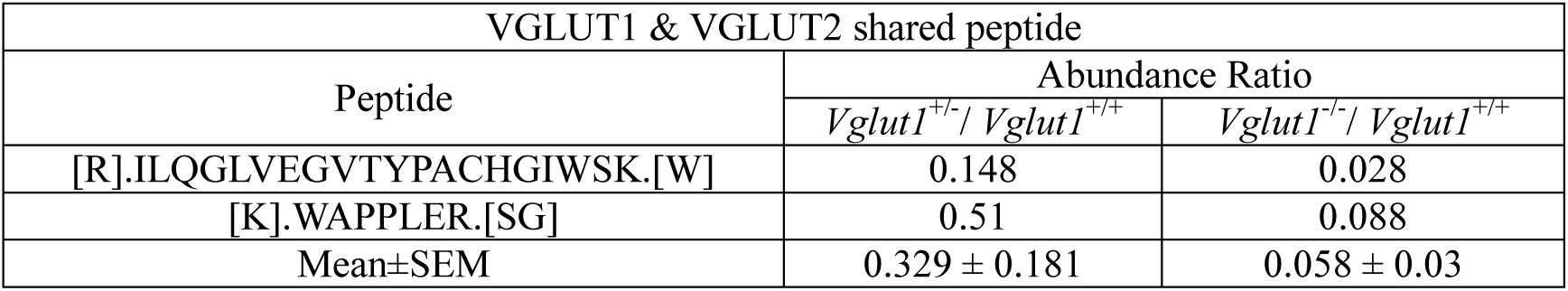

## References

Balschun D, Moechars D, Callaerts-Vegh Z, Vermaercke B, Van Acker N, Andries L, D’Hooge R (2010) Vesicular glutamate transporter VGLUT1 has a role in hippocampal long-term potentiation and spatial reversal learning. Cereb Cortex 20:684–693.

Banerjee S, Vernon S, Jiao W, Choi BJ, Ruchti E, Asadzadeh J, Burri O, Stowers RS, McCabe BD (2021) Miniature neurotransmission is required to maintain Drosophila synaptic structures during ageing. Nat Commun 12:4399.

Belloch F de B, Cortés-Erice M, Herzog E, Zhang XM, Díaz-Perdigon T, Puerta E, Tordera RM (2023) Fast antidepressant action of ketamine in mouse models requires normal VGLUT1 levels from prefrontal cortex neurons. Prog Neuropsychopharmacol Biol Psychiatry 121:110640.

Berardozzi R, Adam V, Martins A, Bourgeois D (2016) Arginine 66 Controls Dark-State Formation in Green-to-Red Photoconvertible Fluorescent Proteins. J Am Chem Soc 138:558–565.

Blumenstock S, Angelo MF, Peters F, Dorostkar MM, Ruf VC, Luckner M, Crux S, Slapakova L, Arzberger T, Claverol S, Herzog E, Herms J (2019) Early defects in translation elongation factor 1α levels at excitatory synapses in α-synucleinopathy. Acta Neuropathol 138:971–986.

Bonanomi D, Benfenati F, Valtorta F (2006) Protein sorting in the synaptic vesicle life cycle. Prog Neurobiol 80:177–217.

Bourgeois D (2023) Single molecule imaging simulations with advanced fluorophore photophysics. Commun Biol 6:53.

Broadhead MJ, Horrocks MH, Zhu F, Muresan L, Benavides-Piccione R, DeFelipe J, Fricker D, Kopanitsa M V, Duncan RR, Klenerman D, Komiyama NH, Lee SF, Grant SGN (2016) PSD95 nanoclusters are postsynaptic building blocks in hippocampus circuits. Sci Rep 6:24626.

Caudle WM, Richardson JR, Wang MZ, Taylor TN, Guillot TS, McCormack AL, Colebrooke RE, Di Monte DA, Emson PC, Miller GW (2007) Reduced vesicular storage of dopamine causes progressive nigrostriatal neurodegeneration. J Neurosci 27:8138–8148.

Cijsouw T, Weber JP, Broeke JH, Broek JAC, Schut D, Kroon T, Saarloos I, Verhage M, Toonen RF (2014) Munc18-1 redistributes in nerve terminals in an activity- and PKC-dependent manner. J Cell Biol 204:759–775.

Daniels RW, Collins CA, Chen K, Gelfand M V, Featherstone DE, DiAntonio A (2006) A single vesicular glutamate transporter is sufficient to fill a synaptic vesicle. Neuron 49:11–16.

Daniels RW, Collins CA, Gelfand M V, Dant J, Brooks ES, Krantz DE, DiAntonio A (2004) Increased expression of the Drosophila vesicular glutamate transporter leads to excess glutamate release and a compensatory decrease in quantal content. J Neurosci 24:10466–10474.

Darcy KJ, Staras K, Collinson LM, Goda Y (2006) Constitutive sharing of recycling synaptic vesicles between presynaptic boutons. Nat Neurosci 9:315–321.

Denker A, Rizzoli SO (2010) Synaptic vesicle pools: an update. Front Synaptic Neurosci 2:135.

Deschout H, Cella Zanacchi F, Mlodzianoski M, Diaspro A, Bewersdorf J, Hess ST, Braeckmans K (2014) Precisely and accurately localizing single emitters in fluorescence microscopy. Nat Methods 11:253–266.

Edwards RH (2007) The neurotransmitter cycle and quantal size. Neuron 55:835–858.

Foss SM, Li H, Santos MS, Edwards RH, Voglmaier SM (2013) Multiple dileucine-like motifs direct VGLUT1 trafficking. J Neurosci 33:10647–10660.

Fremeau RT, Kam K, Qureshi T, Johnson J, Copenhagen DR, Storm-Mathisen J, Chaudhry FA, Nicoll RA, Edwards RH (2004) Vesicular glutamate transporters 1 and 2 target to functionally distinct synaptic release sites. Science 304:1815–1819.

Garcia-Garcia AL, Elizalde N, Matrov D, Harro J, Wojcik SM, Venzala E, Ramírez MJ, Del Rio J, Tordera RM (2009) Increased vulnerability to depressive-like behavior of mice with decreased expression of VGLUT1. Biol Psychiatry 66:275–282.

Herzog E, Bellenchi GC, Gras C, Bernard V, Ravassard P, Bedet C, Gasnier B, Giros B, El Mestikawy S (2001) The existence of a second vesicular glutamate transporter specifies subpopulations of glutamatergic neurons. J Neurosci 21:RC181.

Herzog E, Nadrigny F, Silm K, Biesemann C, Helling I, Bersot T, Steffens H, Schwartzmann R, Nagerl U V., El Mestikawy S, Rhee J, Kirchhoff F, Brose N (2011) In Vivo Imaging of Intersynaptic Vesicle Exchange Using VGLUT1Venus Knock-In Mice. Journal of Neuroscience 31:15544–15559.

Herzog E, Takamori S, Jahn R, Brose N, Wojcik SM (2006) Synaptic and vesicular co-localization of the glutamate transporters VGLUT1 and VGLUT2 in the mouse hippocampus. J Neurochem 99:1011–1018.

Izeddin I, Boulanger J, Racine V, Specht CG, Kechkar A, Nair D, Triller A, Choquet D, Dahan M, Sibarita JB (2012) Wavelet analysis for single molecule localization microscopy. Opt Express 20:2081–2095.

Käll L, Canterbury JD, Weston J, Noble WS, MacCoss MJ (2007) Semi-supervised learning for peptide identification from shotgun proteomics datasets. Nat Methods 4:923–925.

Kalla S, Stern M, Basu J, Varoqueaux F, Reim K, Rosenmund C, Ziv NE, Brose N (2006) Molecular dynamics of a presynaptic active zone protein studied in Munc13-1-enhanced yellow fluorescent protein knock-in mutant mice. Journal of Neuroscience 26:13054–13066.

Kavalali ET (2015) The mechanisms and functions of spontaneous neurotransmitter release. Nat Rev Neurosci 16:5–16.

Kechkar A, Nair D, Heilemann M, Choquet D, Sibarita J-B (2013) Real-time analysis and visualization for single-molecule based super-resolution microscopy. PLoS One 8:e62918.

Lakso M, Pichel JG, Gorman JR, Sauer B, Okamoto Y, Lee E, Alt FW, Westphal H (1996) Efficient in vivo manipulation of mouse genomic sequences at the zygote stage. Proc Natl Acad Sci U S A 93:5860–5865.

Lohr KM, Bernstein AI, Stout KA, Dunn AR, Lazo CR, Alter SP, Wang M, Li Y, Fan X, Hess EJ, Yi H, Vecchio LM, Goldstein DS, Guillot TS, Salahpour A, Miller GW (2014) Increased vesicular monoamine transporter enhances dopamine release and opposes Parkinson disease-related neurodegeneration in vivo. Proc Natl Acad Sci U S A 111:9977–9982.

McKinney SA, Murphy CS, Hazelwood KL, Davidson MW, Looger LL (2009) A bright and photostable photoconvertible fluorescent protein. Nat Methods 6:131–133.

Minerbi A, Kahana R, Goldfeld L, Kaufman M, Marom S, Ziv NE (2009) Long-term relationships between synaptic tenacity, synaptic remodeling, and network activity. PLoS Biol 7:e1000136.

Miyazaki T, Fukaya M, Shimizu H, Watanabe M (2003) Subtype switching of vesicular glutamate transporters at parallel fibre-Purkinje cell synapses in developing mouse cerebellum. Eur J Neurosci 17:2563–2572.

Moechars D, Weston MC, Leo S, Callaerts-Vegh Z, Goris I, Daneels G, Buist A, Cik M, van der Spek P, Kass S, Meert T, D’Hooge R, Rosenmund C, Hampson RM (2006) Vesicular glutamate transporter VGLUT2 expression levels control quantal size and neuropathic pain. J Neurosci 26:12055–12066.

Mutch SA, Kensel-Hammes P, Gadd JC, Fujimoto BS, Allen RW, Schiro PG, Lorenz RM, Kuyper CL, Kuo JS, Bajjalieh SM, Chiu DT (2011) Protein quantification at the single vesicle level reveals that a subset of synaptic vesicle proteins are trafficked with high precision. J Neurosci 31:1461–1470.

Nagai T, Ibata K, Park ES, Kubota M, Mikoshiba K, Miyawaki A (2002) A variant of yellow fluorescent protein with fast and efficient maturation for cell-biological applications. Nat Biotechnol 20:87–90.

Nakakubo Y, Abe S, Yoshida T, Takami C, Isa M, Wojcik SM, Brose N, Takamori S, Hori T (2020) Vesicular Glutamate Transporter Expression Ensures High-Fidelity Synaptic Transmission at the Calyx of Held Synapses. Cell Rep 32:108040.

Nienhaus K, Nienhaus GU, Wiedenmann J, Nar H (2005) Structural basis for photo-induced protein cleavage and green-to-red conversion of fluorescent protein EosFP. Proc Natl Acad Sci U S A 102:9156–9159.

Paez-Segala MG, Sun MG, Shtengel G, Viswanathan S, Baird MA, Macklin JJ, Patel R, Allen JR, Howe ES, Piszczek G, Hess HF, Davidson MW, Wang Y, Looger LL (2015) Fixation-resistant photoactivatable fluorescent proteins for CLEM. Nat Methods 12:215–218, 4 p following 218.

Pan P-Y, Marrs J, Ryan TA (2015) Vesicular glutamate transporter 1 orchestrates recruitment of other synaptic vesicle cargo proteins during synaptic vesicle recycling. J Biol Chem 290:22593–22601.

Penn AC, Zhang CL, Georges F, Royer L, Breillat C, Hosy E, Petersen JD, Humeau Y, Choquet D (2017) Hippocampal LTP and contextual learning require surface diffusion of AMPA receptors. Nature 549:384–388.

Racine V, Sachse M, Salamero J, Fraisier V, Trubuil A, Sibarita J-B (2007) Visualization and quantification of vesicle trafficking on a three-dimensional cytoskeleton network in living cells. J Microsc 225:214–228.

Rizzoli SO, Betz WJ (2004) The structural organization of the readily releasable pool of synaptic vesicles. Science 303:2037–2039.

Rizzoli SO, Betz WJ (2005) Synaptic vesicle pools. Nat Rev Neurosci 6:57–69 Available at: http://www.ncbi.nlm.nih.gov/pubmed/15611727.

Rothman JS, Kocsis L, Herzog E, Nusser Z, Silver RA (2016) Physical determinants of vesicle mobility and supply at a central synapse. Elife 5.

Sakaba T, Schneggenburger R, Neher E (2002) Estimation of quantal parameters at the calyx of Held synapse. Neurosci Res 44:343–356.

Shroff H, Galbraith CG, Galbraith JA, Betzig E (2008) Live-cell photoactivated localization microscopy of nanoscale adhesion dynamics. Nat Methods 5:417–423.

Siksou L, Silm K, Biesemann C, Nehring RB, Wojcik SM, Triller A, El Mestikawy S, Marty S, Herzog E (2013) A role for vesicular glutamate transporter 1 in synaptic vesicle clustering and mobility. European Journal of Neuroscience 37:1631–1642.

Song H, Ming G, Fon E, Bellocchio E, Edwards RH, Poo M (1997) Expression of a putative vesicular acetylcholine transporter facilitates quantal transmitter packaging. Neuron 18:815–826.

Staras K, Branco T, Burden JJ, Pozo K, Darcy K, Marra V, Ratnayaka A, Goda Y (2010) A vesicle superpool spans multiple presynaptic terminals in hippocampal neurons. Neuron 66:37–44.

Subach F V, Patterson GH, Manley S, Gillette JM, Lippincott-Schwartz J, Verkhusha V V (2009) Photoactivatable mCherry for high-resolution two-color fluorescence microscopy. Nat Methods 6:153–159.

Sudhof TC (2004) The synaptic vesicle cycle. Annu Rev Neurosci 27:509–547.

Sutton MA, Ito HT, Cressy P, Kempf C, Woo JC, Schuman EM (2006) Miniature neurotransmission stabilizes synaptic function via tonic suppression of local dendritic protein synthesis. Cell 125:785–799.

Takamori S et al. (2006) Molecular anatomy of a trafficking organelle. Cell 127:831–846.

Takamori S (2016) Presynaptic Molecular Determinants of Quantal Size. Front Synaptic Neurosci 8:2.

Taylor TN, Alter SP, Wang M, Goldstein DS, Miller GW (2014) Reduced vesicular storage of catecholamines causes progressive degeneration in the locus ceruleus. Neuropharmacology 76 Pt A:97–105.

Thédié D, Berardozzi R, Adam V, Bourgeois D (2017) Photoswitching of Green mEos2 by Intense 561 nm Light Perturbs Efficient Green-to-Red Photoconversion in Localization Microscopy. J Phys Chem Lett 8:4424–4430.

Tsuriel S, Geva R, Zamorano P, Dresbach T, Boeckers T, Gundelfinger ED, Garner CC, Ziv NE (2006) Local sharing as a predominant determinant of synaptic matrix molecular dynamics. PLoS Biol 4:e271.

Turrigiano G (2012) Homeostatic Synaptic Plasticity: Local and Global Mechanisms for Stabilizing Neuronal Function. Cold Spring Harb Perspect Biol 4:a005736–a005736.

Walmsley B, Edwards FR, Tracey DJ (1988) Nonuniform release probabilities underlie quantal synaptic transmission at a mammalian excitatory central synapse. J Neurophysiol 60:889–908.

Watanabe S, Boucrot E (2017) Fast and ultrafast endocytosis. Curr Opin Cell Biol 47:64–71.

Watanabe S, Trimbuch T, Camacho-Pérez M, Rost BR, Brokowski B, Söhl-Kielczynski B, Felies A, Davis MW, Rosenmund C, Jorgensen EM (2014) Clathrin regenerates synaptic vesicles from endosomes. Nature 515:228–233.

Watson ET, Pauers MM, Seibert MJ, Vevea JD, Chapman ER (2023) Synaptic vesicle proteins are selectively delivered to axons in mammalian neurons. Elife 12.

Weston MC, Nehring RB, Wojcik SM, Rosenmund C (2011) Interplay between VGLUT isoforms and endophilin A1 regulates neurotransmitter release and short-term plasticity. Neuron 69:1147–1159.

Westphal V, Rizzoli SO, Lauterbach MA, Kamin D, Jahn R, Hell SW (2008) Video-rate far-field optical nanoscopy dissects synaptic vesicle movement. Science 320:246–249.

Wilhelm BG, Mandad S, Truckenbrodt S, Kröhnert K, Schäfer C, Rammner B, Koo SJ, Claßen GA, Krauss M, Haucke V, Urlaub H, Rizzoli SO (2014) Composition of isolated synaptic boutons reveals the amounts of vesicle trafficking proteins. Science 344:1023–1028.

Wojcik SM, Rhee JS, Herzog E, Sigler A, Jahn R, Takamori S, Brose N, Rosenmund C (2004) An essential role for vesicular glutamate transporter 1 (VGLUT1) in postnatal development and control of quantal size. Proc Natl Acad Sci U S A 101:7158–7163.

Wulffele J, Thédié D, Glushonkov O, Bourgeois D (2022) mEos4b Photoconversion Efficiency Depends on Laser Illumination Conditions Used in PALM. J Phys Chem Lett 13:5075–5080.

Zhang M, Chang H, Zhang Y, Yu J, Wu L, Ji W, Chen J, Liu B, Lu J, Liu Y, Zhang J, Xu P, Xu T (2012) Rational design of true monomeric and bright photoactivatable fluorescent proteins. Nat Methods 9:727–729.

Zhang XM, François U, Silm K, Angelo MF, Fernandez-Busch MV, Maged M, Martin C, Bernard V, Cordelières FP, Deshors M, Pons S, Maskos U, Bemelmans AP, Wojcik SM, El Mestikawy S, Humeau Y, Herzog E (2019) A proline-rich motif on VGLUT1 reduces synaptic vesicle super-pool and spontaneous release frequency. Elife 8:1–24.

